# Cannabidiol exerts anti-inflammatory effects but maintains T effector memory cell differentiation: A Single-Cell Study in Humans

**DOI:** 10.1101/2025.07.30.667742

**Authors:** Debora L. Gisch, Sachiko Koyama, Jumar Etkins, Gerald C So, Daniel J. Fehrenbach, Jessica Bo Li Lu, Ying-Hua Cheng, Ricardo Melo Ferreira, Evan Rajadhyaksha, Kelsey McClara, Mahla Asghari, Asif A. Sharfuddin, Pierre C. Dagher, Laura M. Snell, Meena S Madhur, Rafael B. Polidoro, Zeruesenay Desta, Michael T. Eadon

## Abstract

Cannabidiol is widely available and often used for pain management. Individuals with kidney disease or renal allografts have limited analgesia options. We conducted a Phase 1 human study to compare the peripheral immune cell distribution before (pre-cannabidiol) and after exposure to cannabidiol at steady state (post-cannabidiol). This *ex vivo* study included specimens from 23 participants who received oral cannabidiol (up to 5 mg/kg twice daily) for 11 days. Lymphocytes were isolated and stimulated with anti-CD3/CD28 antibodies, with or without tacrolimus. Pharmacodynamic responses were assessed via CellTiter-Glo® proliferation, scRNA-seq, cytokine assays, and flow cytometry. Steady-state plasma concentrations of CBD were quantified via tandem mass spectrometry. We identified an increased proportion of T effector memory (TEM) cells post-cannabidiol (22% increase, *P-value* of 3.2 x 10^-32^), which correlated with CBD plasma concentrations (*Pearson Corr= 0.77, P-value < 0.01*). Post-cannabidiol cytokine assays revealed elevated proinflammatory IL-6 protein levels and anti-inflammatory IL-10 levels (*adjusted P-values < 0.0001*). Cannabidiol reduced overall T and B lymphocyte proliferation with additive immunosuppressive effects to tacrolimus. In flow cytometry, the proportion of TEM and TEMRA increased post-cannabidiol with tacrolimus (*P-values < 0.05*). Cannabidiol exhibits mixed immunomodulatory effects with pro- and anti-inflammatory signals. Understanding the clinical safety of cannabidiol use is important given the paucity of pain control options available for immunocompromised transplant populations.

## Introduction

Cannabidiol (CBD) is approved by the United States Food and Drug Administration (FDA) for certain seizure disorders^1^. CBD is also widely used off-label for the management of chronic pain.^2^ Individuals with chronic kidney disease (CKD) or those who are transplant allograft recipients face challenges with analgesia because non-steroidal anti-inflammatory drugs (NSAIDs) are nephrotoxic and pose a significant risk for kidney function decline. CBD is a particularly attractive alternative for off-label pain control in these populations^3^. However, special considerations are required for allograft recipients who often receive calcineurin-inhibitor (CNI) based immunosuppressive regimens. Allograft recipients are at risk for pharmacokinetic drug-drug interactions (DDIs) with tacrolimus (TAC) and everolimus^4–6^. However, less is understood regarding the interactions between CBD and CNIs within the immune system. This warrants particular attention because CBD and CNIs are involved in the deregulation of nuclear factor of activated T-cells (NFAT)^7^.

Allograft recipients require lifelong immunosuppression, placing them at risk for opportunistic infections.^8^ The FDA-approved form of cannabidiol (Epidiolex) contains a warning regarding the increased risk of infection, without a mechanistic explanation for this adverse event^9^. The immunomodulatory properties of CBD have been extensively studied in cell culture systems. For example, CBD has been shown to reduce T cell proliferation and proinflammatory cytokine secretion, in part through the suppression of NFAT and Activator Protein-1 (AP-1)^7,10,11^. Most literature supports the anti-inflammatory effect of CBD, but some studies also reveal a mixed picture of pro- and anti-inflammatory effects^12^. However, clinical studies addressing the impact of CBD on the immune system are lacking. Thus, we sought to understand the effects of CBD on the immune system in healthy participants before and after two weeks of CBD treatment.

In this study, we assessed the pharmacodynamic effects of CBD and drug-drug interaction (DDI) between CBD and tacrolimus on the immune system in human participants. Participants took 14 days of controlled Epidiolex administration at 5 mg/kg by mouth twice daily. An enriched set of their lymphocytes underwent single-cell RNA sequencing, proliferation assays, cytokine assessment, and flow cytometry before (pre-CBD) and after (post-CBD) 11 days of exposure to assess the comprehensive changes in the lymphocyte landscape. Pre-CBD and post-CBD blood samples were divided and sequenced with (CD3/CD28) and without anti-CD3/CD28 (no CD3/CD28) antibody stimulation to simulate antigen exposure, and after stimulation with the addition of a defined concentration of tacrolimus (5 ng/ml) to assess the pharmacodynamic DDI (CD3/CD28 +TAC). We hypothesized that CBD would exert anti-inflammatory effects and reduce immune cell proliferation. This hypothesis was only partially supported as CBD led to a mixed set of pro- and anti-inflammatory effects in the human participants. Some effects were synergistic, and others were antagonistic to tacrolimus-mediated ones. The immunomodulatory effects of CBD included significant concentration-dependent changes in T effector memory cell (TEM) proportions and cell-cell signaling.

## Results

### PARTICIPANTS

A total of 41 participants completed a pharmacokinetic study of CBD, which closed enrollment on 7/15/25. Participants received 11 days of cannabidiol at 5 mg/kg twice daily before steady state pharmacokinetic sampling on day 12. A subset of participants (N=23) provided blood samples for advanced phenotyping of their immune system before CBD exposure (pre-CBD) and at steady-state trough (post-CBD), just before the morning dose on day 12, to determine the effects of CBD on lymphocyte distribution, expression, and signaling. The peripheral blood samples underwent an experimental protocol to enrich the lymphocyte population.^13^ Advanced phenotyping included pre-CBD single-cell/nucleus RNA-sequencing (scRNA-seq, N = 12), post-CBD scRNA-seq (N = 10 with paired pre-CBD scRNA-seq and pharmacokinetics), CellTiter-Glo® proliferation assays (N = 19), cytokine assays (N = 3), and flow cytometry (N = 3). **Supplementary Table 1** shows which assays were performed for each participant. A flow diagram (**Figure 1**) illustrates the study enrollment, timeline, and experimental technologies used to interrogate the specimens.

**Figure 1:**
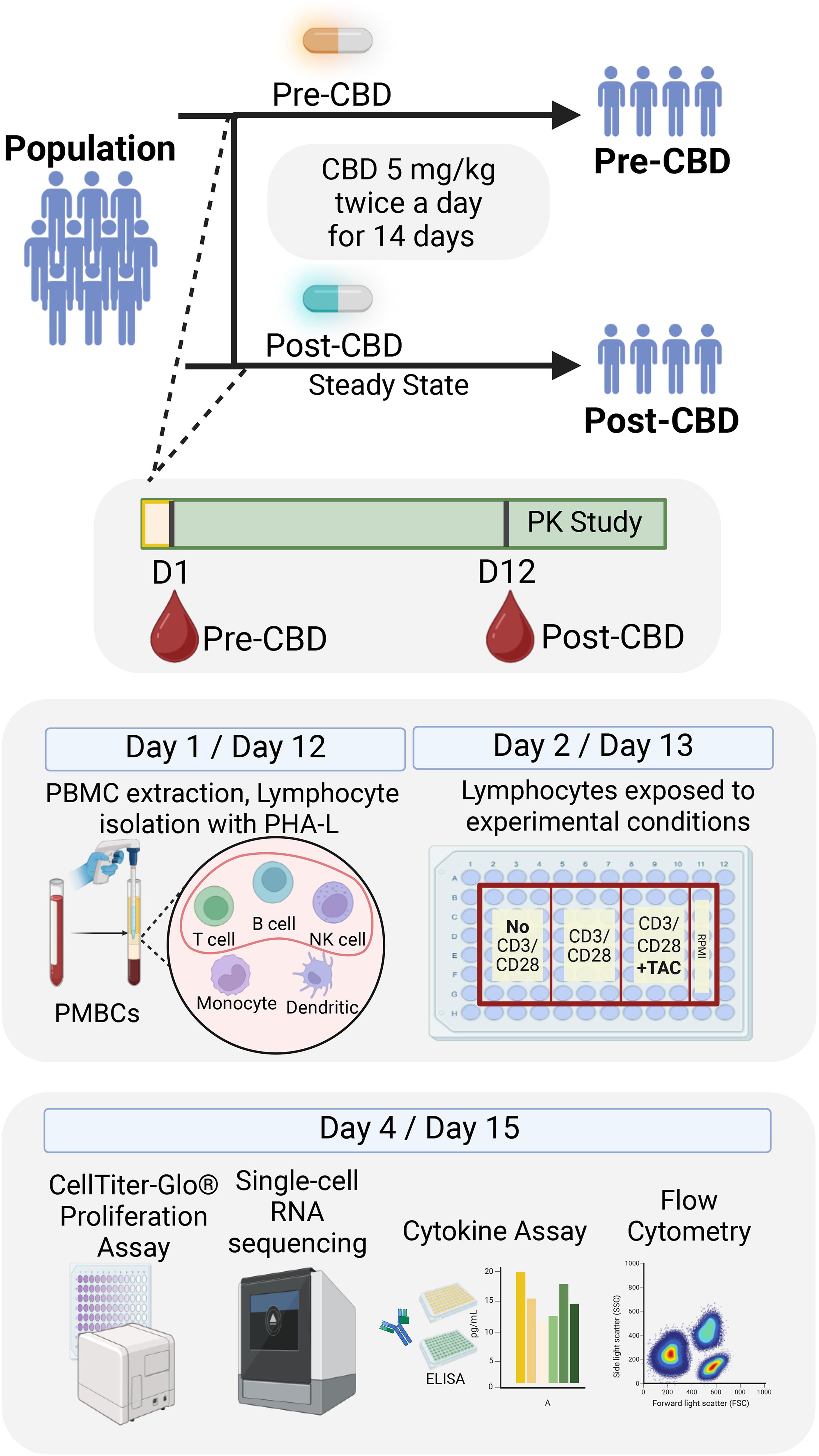
**Operational workflow**. The workflow is provided for the pharmacokinetic study of steady-state cannabidiol (CBD) in healthy human volunteers, highlighting the time points of sample acquisition for molecular analysis. Samples were acquired at two timepoints: pre-CBD or before CBD administration (D1), and post-CBD at steady state (D12). Each participant took oral CBD up to 5 mg/kg twice daily for 14 days. The blood samples underwent 72 hours of processing. On Day 1 / Day 12, lymphocytes were isolated. On Day 2 / Day 13, cells were divided into conditions: baseline without stimulation (No CD3/CD28); CD3/CD28 stimulation (CD3/CD28), and stimulated + tacrolimus (CD3/CD28 + TAC). After 72 hours, cells were interrogated by CellTiter-Glo®, single-cell RNA sequencing (scRNA-seq), cytokine measurement, and flow cytometry. Between D12 and D14, participants underwent pharmacokinetic sampling for 48h to obtain CBD concentrations at steady-state. Created in BioRender. Gisch, D. (2025) https://BioRender.com/416zmh0

### THE EFFECTS OF CBD ON IMMUNE CELL PROLIFERATION AND DISTRIBUTION

Cell proliferation was assessed after anti-CD3/CD28 stimulation (CD3/CD28) at pre- and post-CBD timepoints using the CellTiter-Glo® assay (**Figure 2a**). To assess proliferation, we calculated the ratio of luminescence in the CD3/CD28 to no CD3/CD28 conditions in both pre-CBD and post-CBD. Pre-CBD, the CD3/CD28 to No CD3/CD28 ratio was 2.04 ± 0.574. Post-CBD, the ratio was significantly reduced to 1.5 ± 0.571 (*P-value = 0.0085*), consistent with a 54% reduction in cell proliferation from individuals treated with CBD for 11 days (**Supplementary Table 2**). To understand the cellular heterogeneity underlying the differences observed in the cell proliferation assay, we performed scRNA-seq on the no CD3/CD28 and CD3/CD28 conditions at the pre-CBD and post-CBD timepoints. After quality control, the scRNA-seq dataset consisted of 314,746 cells across four conditions (**Figure 2b**): No CD3/CD28 -CBD (91,865 cells), No CD3/CD28 +CBD (54,914 cells), CD3/CD28 -CBD (108,423 cells), and CD3/CD28 +CBD (59,560 cells).

**Figure 2:**
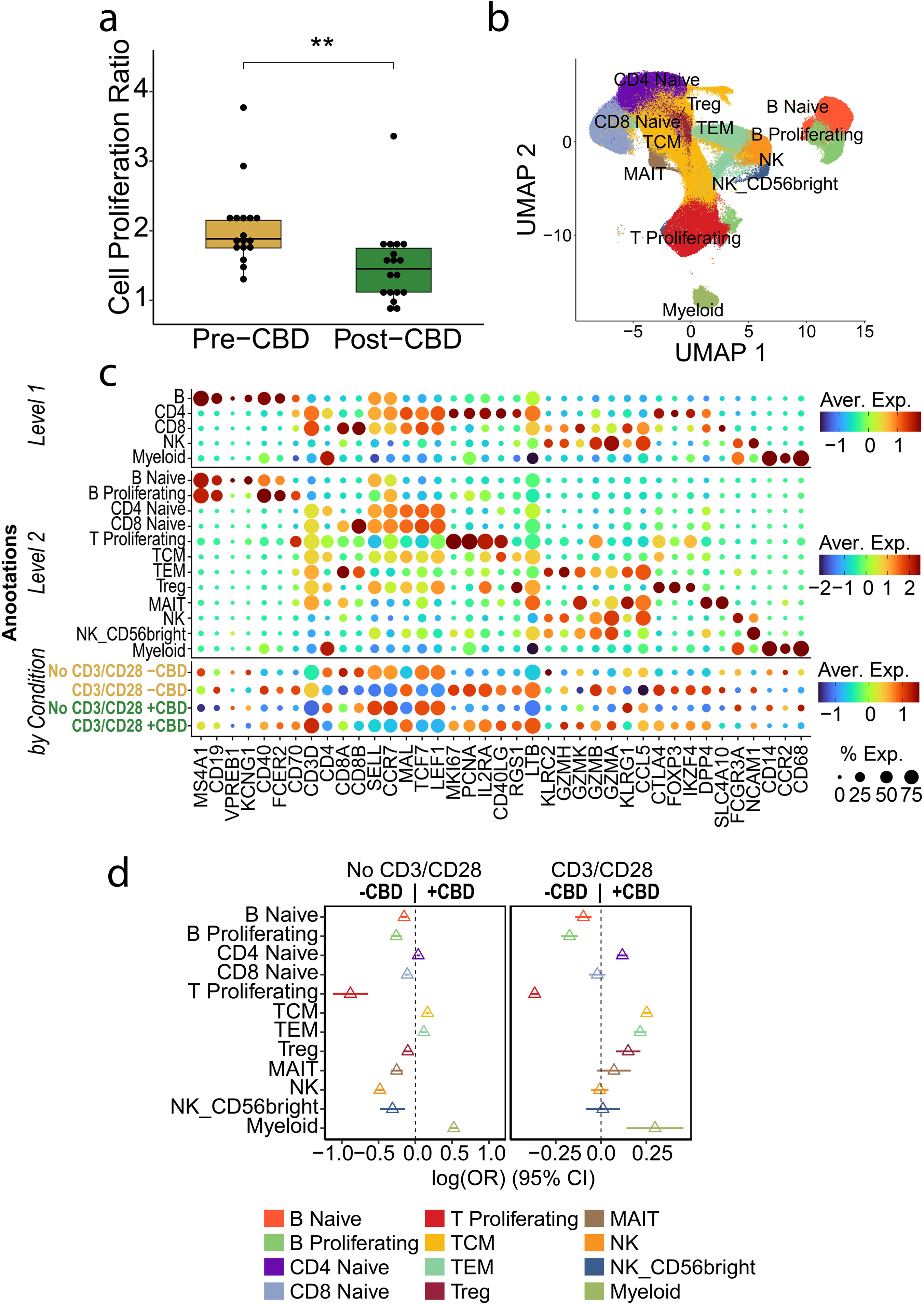
Molecular characterization of lymphocytes. Lymphocyte proliferation, annotation, and distribution were determined at pre- and post-CBD timepoints. (a) The CellTiter-Glo® assay assessed cell viability at the pre-CBD (N=17) and post-CBD (N=18) timepoints. The proliferation ratio is the measured values of the stimulated (CD3/CD28) condition normalized to the no CD3/CD28 condition (No CD3/CD28). Cell proliferation was reduced post-CBD (*P-value < 0.001 by t-test*). (b) UMAP with annotations from scRNA-seq (N=59) experiments at the pre-CBD and post-CBD timepoints for the CD3/CD28 and No CD3/CD28 conditions (321,327 cells displayed). Twelve cell type clusters were annotated: B naive, B Proliferating, CD4 Naive, CD8 Naïve, T Proliferating, T Central Memory (**TCM**), T effector memory (**TEM**), T regulatory (**Treg**), Mucosal-associated invariant T (**MAIT**), Natural Killer (**NK**), Natural Killer CD56+ (**NK_CD56brigh**t), and retained Myeloid. (c) Canonical marker gene expression of peripheral blood mononuclear cell (PBMC) types was used to annotate cell types. Three levels of annotation are provided: Level 1 includes B cells (MS4A1, CD19), CD4+ lymphocytes (CD4), CD8 lymphocytes (CD8A, CD8B), NK cells (NCAM1), and Myeloid cells (CD14, CCR2, and *CD68*). Level 2 annotations increased specificity with 12 cell types characterized by the presence of naïve B *(VPREGB1 and KCNG1), CD4 naïve and CD8 naïve* (*SELL*, *CCR7*, *MAL*, *TCF7* and *LEF1*), B Proliferating (*CD40*, *FCER2* and *CD70*) and T proliferating (*MKI67*, *PCNA* and *IL2RA*) and TCM (CCR7, *SELL, CD40LG*, *RGS1* and *LTB),* TEM (*GZMH*, *GZMK*, *GZMB*, *GZMA*, *KLRG1* and *CCL5*), Treg (*CTLA4*, *FOXP3* and *IKZF4*), MAIT (*DPP4* and *SLC4A10*). The dot size indicates the percentage of cells expressing each gene; color represents the average expression (log-normalized), from low (dark blue) to high (dark red). (d) Cell proportions changed between the pre-CBD and post-CBD timepoints in the No CD3/CD28 and CD3/CD28 conditions. Among other changes, an increase in TEM was observed post-CBD, while a decrease in proliferating B and T lymphocytes was observed. Forest plots display the log_2_ odds ratio between timepoints across the Level 2 annotations. For each cell type, the central triangle marks the estimated difference in proportions between conditions, and the adjoining horizontal line depicts its 95 % confidence interval. A confidence interval that intersects the vertical dashed null line indicates no statistically significant difference, whereas intervals that remain entirely on one side denote significance at *P* -value< 0.05.

We clustered and annotated cells at both low (Level 1) and high (Level 2) resolution based on marker gene expression (**Supplementary Table 3**). The Level 1 annotation consisted of 5 cell types (B, CD4, CD8, NK, and Myeloid) used in downstream cross-cutting differential expression and pathways analyses. To assess granular cell type distribution, we used Level 2 annotations to define 12 cell types (**Figure 2c**). These 12 subclusters represented the following cell types with parenthetical marker genes: naïve B lymphocyte (*CD1*9, *KCNG1*); proliferating B lymphocyte (*CD19*, *CD40*); naïve CD4+ T lymphocyte (*CD4*, *SELL*, *CCR7, TCF7*); naïve CD8+ T lymphocyte (*CD8B*, *SELL*, *CCR7, TCF7*); proliferating T lymphocyte (*CD3D*, *MKI67*, *PCNA*); T central memory or TCM (*CD3D, SELL, RGS1, D40LG*); T effector memory or TEM (*CD3D, KLRC2, GZMH, CCL5* with absence of *CCR7* and *SELL*); T regulatory lymphocyte or Treg (*FOXP3*, *CTLA4,* and IKZF4); Mucosal-Associated Invariant T or MAIT (*DPP4*, *C4A10)*; Natural Killer or NK (*FCGR3A*); Natural Killer *CD56* positive orNK_CD56bright) (*NCAM1*); and Myeloid (*CD14, CCR2, CD68*) – only a limited population of myeloid cells were retained due to our experimental selection for lymphocyte populations. Further sub-clustering of the proliferating T lymphocyte, TCM, and TEM clusters into *CD4*+ and *CD8*+ T lymphocytes was not feasible because they clustered based on their proliferative or memory features rather than *CD4*+ and *CD8*+ expression (**Supplementary** Figure 1). However, a greater proportion of TCMs were *CD4+*, while a greater proportion of TEMs were *CD8+*.

The immune cell distribution of participants changed significantly after 11 days of CBD administration (**Figure 2d, Supplementary Table 4**). In the no CD3/CD28 baseline condition, the proportion of proliferating T and B lymphocytes, naïve B lymphocytes, MAITs, and NK cells was reduced post-CBD compared to pre-CBD. Although the absolute quantity of all cell types was reduced post-CBD, we observed significant *relative increases* in TCMs and TEMs when comparing the same conditions. With CD3/CD28 stimulation, we again observed reductions in proliferating T (36% reduction, 95% CI: -39 to -34), B (17% reduction, 95% CI: -22 to - 13) lymphocytes, and naïve B lymphocytes post-CBD. A greater proportion of TCMs (25% increase, 95% CI: 23 to 27) and TEMs (21% increase, 95% CI: 18 to 25) were also observed post-CBD. One key difference between the CD3/CD28 and no CD3/CD28 conditions was the change in Treg proportion, which increased post-CBD with CD3/CD28. Myeloid cell populations were upregulated post-CBD, but we interpret these results with caution given our experimental design. Finally, we identified changes in expression patterns across all Level 1 and Level 2 resolution cell types in the CD3/CD28 and no CD3/CD28 conditions between participants pre-CBD and post-CBD (**Supplementary** Figure 2**, Supplementary Table 5**). *CD69*, an early activation receptor, was downregulated in naïve T cells post-CBD. *KIT*, a type III receptor tyrosine kinase involved in hematopoiesis, was downregulated in proliferative T cells post-CBD, but was not differentially expressed in the Treg population.

### CBD INCREASES T EFFECTOR MEMORY CELL PROPORTION IN A CONCENTRATION-DEPENDENT MANNER

A group of participants (N = 10) had both pharmacokinetic and scRNA-seq data post-CBD (**Figure 3a**). The study population had a median age of 42 ± 18.5 years, and 80% were female. Most individuals self-identified as white or European American (70%). Plasma concentrations of CBD were assessed for the steady state phase of CBD administration on day 12 at 8 sampling time points over a 12-hour dosing interval (**Figure 3b**). Pharmacokinetic parameters were derived by non-compartmental analysis (**Figure 3c**). The mean CBD maximal concentration (C_max,ss_) was 438.0 ng/ml (95% CI: 263.5 to 612.5 ng/mL). The mean trough concentration (C_0h,ss_) was 141.5 ng/mL, with substantial variability of 95% CI: 64.2 to 218.8 ng/mL. Because we had identified an increase in TEMs post-CBD, we tested whether the TEM proportion correlated with either C_0h,ss_ or C_max,ss_. The strongest correlation between concentration and TEM proportion was seen at the C_0h,ss_ timepoint (*correlation R = 0.77, P-value = 0.01*), when the scRNA-seq data were acquired (**Figure 3d**). We identified a trend toward a negative correlation between proliferative T cells and increasing CBD concentrations at multiple time points. Because the absolute quantities of all cell types were reduced post-CBD, the increased TEM proportion is due to a relatively greater reduction in non-TEM cell abundance post-CBD. We did not identify a significant positive or negative correlation with the Treg proportion.

**Figure 3:**
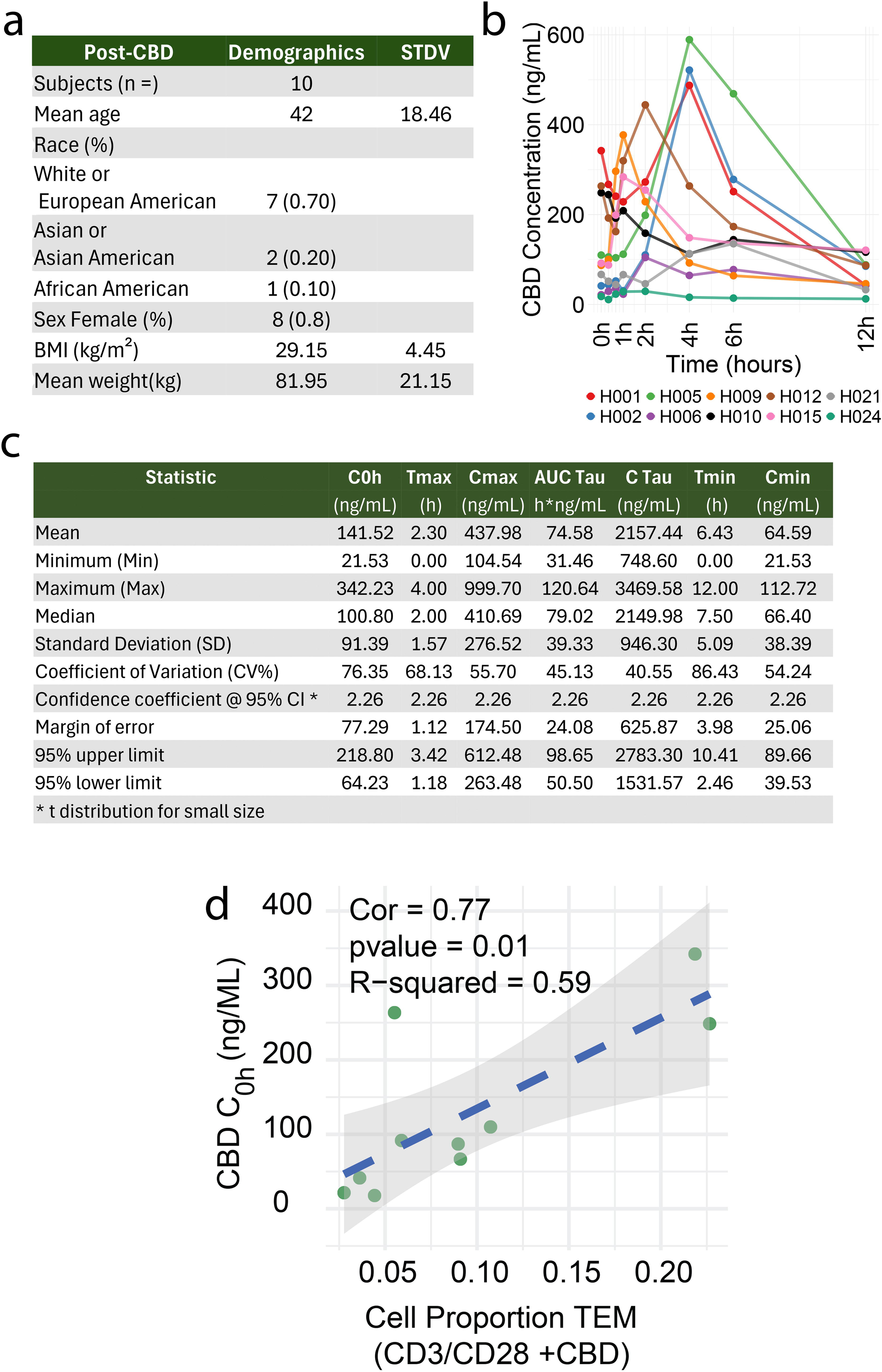
CBD concentration-dependent effects on cell proportion. In the post-CBD phase, 10 participants had overlapping sc/snRNA-seq and pharmacokinetic data. (a) Demographics of the participants following CBD treatment. The mean age of the participants is 42 ± 18.5 years. (b) Concentration to time curves of CBD concentration levels (ng/mL) at time points starting on Day 12: 0h, 0.33h, 0.66h, 1h, 2h, 4h, 6h, and 12h while at CBD steady-state. Some inter-individual variability was observed in the pharmacokinetic profiles of participants. (c) Pharmacokinetic parameters of CBD include the through concentration (C_0h_), time to reach maximum concentration (T_max_), maximum concentration (C_max_), area under the curve (AUC_0-tau_), concentration at the end of the dosing interval (C_Tau_), time to reach minimum concentration (T_min_), and minimum concentration (C_min_). The values are presented as mean, minimum (Min), maximum (Max), median, standard deviation (SD), coefficient of variation (CV%), confidence coefficient at 95% confidence interval (CI), margin of error, and the 95% upper and lower limits. (d) A scatter plot shows the relationships between immune cell proportions and CBD concentrations in participants. The dark green dots represent the proportion of TEM cells versus CBD trough concentration at steady state (C_0h_), with a Pearson correlation coefficient (Cor) of 77%, a P-value of 0.01, and an R-squared value of 0.59. The dashed lines indicate the linear regression fits, performed using the Pearson correlation by lm method in ggplot2 (v.3.5.1).

### CBD MODULATES INTERCELLULAR COMMUNICATION AND PATHWAYS OF T EFFECTOR MEMORY CELLS

To investigate the mechanism of how TEMs respond to CBD, we profiled cell–cell communication comparing pre-CBD and post-CBD receptor-ligand (R-L) pairs. With TEM as the source, we identified 9 R-L pairs turned on post-CBD and 13 R-L pairs turned off post-CBD (**Figure 4a**). Lymphotoxin-alpha (LTA or Tumor Necrosis Factor (TNF)-beta signaling was altered in multiple cell types. For example, communication was lost from TEM cells to proliferating B lymphocytes, proliferating T lymphocytes, naïve CD4 cells, and myeloid cells through the R-L pair LTA-TNFRSF14. Signaling through the R-L pairs LTA-TNFRSF1B and LTA-TNFRSF1A was also lost in proliferating T lymphocytes.

**Figure 4:**
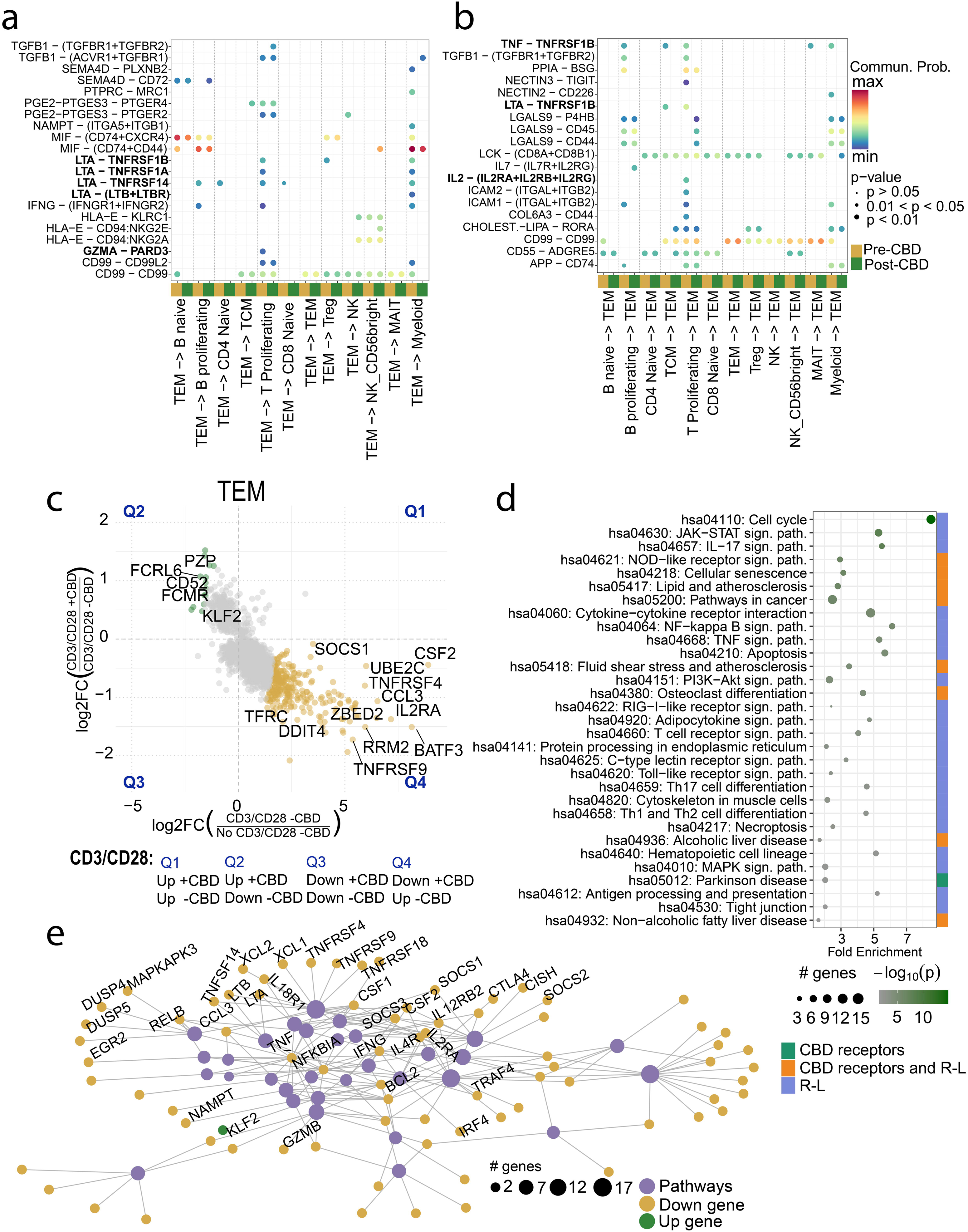
Receptor-ligand interactions and expression signature of T effector memory cells. The receptor-ligand interactions, expression signature, and pathways of T effector memory (TEM) cells were explored pre-CBD and post-CBD. (a) Outgoing signals from TEM cells (source) to all other immune cell types (targets) are displayed in the dotplot, both pre-CBD (dark yellow) and post-CBD (dark green). Directional ligand–receptor (L–R) signaling interactions involving TEM cells change before and after CBD titration. Communication was lost from TEM cells to proliferating B lymphocytes, proliferating T lymphocytes, and naïve CD4 cells through the *LTA-TNFRSF14* R-L, *LTA-TNFRSF1B*, and *LTA-TNFRSF1A* R-L signals. (b) Incoming signals to TEM cells (target) from all other cell types (sources) are displayed. *LTA-TNFRSF1B* signaling was lost in proliferating T lymphocytes and T central memory cells due to a loss of *TNFRSF1B* expression in TEMs. Signaling through TNF to *TNFRSF1B* was lost in proliferating B, T, and TCM lymphocytes. Interleukin-2 (IL-2) signaling through IL-2 receptors was lost between proliferative T lymphocytes and the TEM. (a-b) Communication probability was compared between pre- and post-CBD conditions. The bubble size represents the significance of the L–R pair (*P-value, t-test*). The communication probability is indicated by colormap maxima (Red) and minima (Blue). Axis labels indicate interacting cell types, with TEM cells consistently involved as a source or target in each panel. The set R-L on the plot was filtered to display R-L pairs activated or inhibited by the CBD condition. (c) Gene expression in TEM cells changed across comparisons: *No CD3/CD28 -CBD vs CD3/CD28 -CBD* (x-axis) and *CD3/CD28 +CBD vs CD3/CD28 -CBD* (y-axis). Scatter plot depicts the log₂ fold changes (log₂FC) in gene expression for TEM cells in each comparison. Each point represents a gene with statistically significant differential expression (*adjusted P-value < 0.05*) in both comparisons. Genes exhibiting opposing directions of regulation—up in one comparison and down in the other—are highlighted (green: upregulated with CBD under stimulation; gold: downregulated with CBD under stimulation). (d) The top 300 genes—identified based on discordant regulation across CBD treatment conditions and ranked by Euclidean distance—were analyzed using active-subnetwork enrichment via *pathfindR*. Genes with adjusted P-value < 0.05 were inputed into *pathfindR* using the BioGRID protein-protein interaction network and KEGG gene set. Enriched pathways were subsequently filtered to retain only those containing known cannabinoid (CBD) receptors or receptor–ligand (R–L) interactions, prioritizing biologically-relevant mechanistic processes. The enrichment chart displays the filtered pathways, ranked by -Log₁₀(*P*), representing those most plausibly linked to CBD’s immunomodulatory actions in the context of TEM cell regulation. (e) A graph-based network visualization of enriched KEGG pathways from TEM cells displays the pathways (purple) in Figure D, adding the differential gene expression DEGs on the pathways (post-CBD), *KLF2* was the only up-regulated gene.

In the reverse direction, with TEM as the target cell, we found 8 R-L pairs turned on post-CBD and 11 R-L pairs turned off post-CBD (**Figure 4b**). LTA-TNFRSF1B signaling was again lost in proliferating T lymphocytes, and now in T central memory cells, due to a loss of *TNFRSF1B* expression in TEMs. Analogously, signaling through TNF to TNFRSF1B was lost in proliferating B, T, and TCM lymphocytes. Interleukin-2 (IL-2) signaling through IL-2 receptors was also disrupted between the proliferative T lymphocytes and the TEM. Post-CBD, increased signaling between the Treg and TEM was predicted through LCK-(CD8A+CD8B1) and multiple HLA proteins with either CD8A or CD8B. The complete list of R-L changes between pre-CBD and post-CBD for all 12 cell types is available in **Supplementary Table 6**.

We next made two comparisons of differentially expressed genes (DEGs) to filter the genes most affected by CBD: 1) CD3/CD28 -CBD v. No CD3/CD28 -CBD to remove genes most affected by stimulation and experimental conditions, and 2) CD3/CD28 -CBD v. CD3/CD28 +CBD to then identify genes specifically related to CBD administration regarding TCR stimulation (**Figure 4c**). Genes in quadrant (Q2) were upregulated in the CD3/CD28 +CBD condition. Still, they were downregulated in the No CD3/CD28 -CBD condition, representing an enhanced response in the CD3/CD28 CBD condition relative to both the No CD3/CD28 -CBD and CD3/CD28 -CBD. Conversely, genes in quadrant (Q4) were downregulated in CD3/CD28 CBD and upregulated in CD3/CD28 -CBD, indicating a suppressed response in the CD3/CD28 +CBD condition. This quadrant-based pattern highlights distinct regulatory shifts driven by the CBD (**Supplementary Table 7**). We selected the top 300 genes farthest from the origin for KEGG pathway enrichment analysis to identify 112 significantly enriched pathways (**Supplementary Table 8**).

We then applied an additional literature filter^3^ to prioritize pathways that include known receptors targeted by CBD or genes/proteins involved in R-L interactions previously identified in the cell-cell communication analyses for TEM cells. This integrated approach highlighted a subset of biologically relevant pathways (N=72 with subset in **Figure 4d**). The pathways included cell cycle arrest, JAK-STAT signaling, IL-17 signaling, NOD-like receptor signaling, and cellular senescence, all representing key mechanisms through which CBD modulates the immune cell environment. *PPARG*, a transcription factor and known CBD receptor, is present in the osteoclast differentiation, lipid, atherosclerosis, and non-alcoholic fatty liver disease pathways. The cation channel TRPV2 is also a known interacting protein for CBD and is present in the NOD-like receptor signaling, TRPV4, fluid shear stress, atherosclerosis, and cellular senescence pathways. *TNF* and *IL2RA* were both downregulated in TEMs after CBD administration and members of over 50 pathways enriched in the post-CBD condition. Among DEGs, most were downregulated in the post-CBD condition (**Figure 4e**); however, one of the upregulated genes was *KLF2*, a transcription factor with a purported role in CD8 T cell differentiation and suppression of exhaustion (Supplementary Tables 7, 8, and 9)^14–16^. *KLF2*+ T cells produce IL-10, may be a subpopulation of T regulatory type 1 cells, and possess an anti-inflammatory phenotype.^17^

### PROLIFERATION OF B AND T LYMPHOCYTES

Cellular proliferation in response to CD3/CD28 was reduced post-CBD, as evidenced by the proliferation assay (**Figure 2a**) and scRNA-seq (**Figure 2d**). To explore the mechanism underlying this reduction, we examined cell-cell communication through R-L pairs in proliferating B and T lymphocytes (**Figure 5a**). Three R-L interactions were changed post-CBD: TNF-TNFRSF1B, TGFB1 – (TGFBR1 + TGFBR2), and ICAM1 – (ITGAL+ITGB2). The last R-L pair was turned on (i.e., opposite direction of effect), with TCM cells increasing *ITGAL*+*ITGB2* ligand expression post-CBD (**Supplementary Table 6**). We again assessed the top 300 differentially expressed genes to examine proliferating B and T lymphocyte pathways (**Figure 5b-5c, Supplementary Tables 10, 11, 12, and 13**). Using a parallel analysis to the TEM, we filtered pathways by known CBD receptors and R-L pairs. Proliferating B and T lymphocyte-enriched pathways revealed strong overlaps, including IL-17 signaling, cell cycle, TGF-beta signaling, and JAK-STAT signaling. The sets of genes that overlapped and had the same direction of effect in both cell types were *CCND2*, *HSP90AB1*, *IL2*, *ID2*, *NAMPT*, *DHX33*, *TRAF4*, *HSPD1*, *KSR1*, and *GZMB (*all downregulated), while *RASGRP2* and *CCND2* were upregulated. These DEGs were present in multiple overlapping pathways (**Figure 5d-5e**). Overall, these data support that CBD treatment retards T cell proliferation through reductions in cell-cell signaling (TNF, TGFBR1, and IL-2).

**Figure 5:**
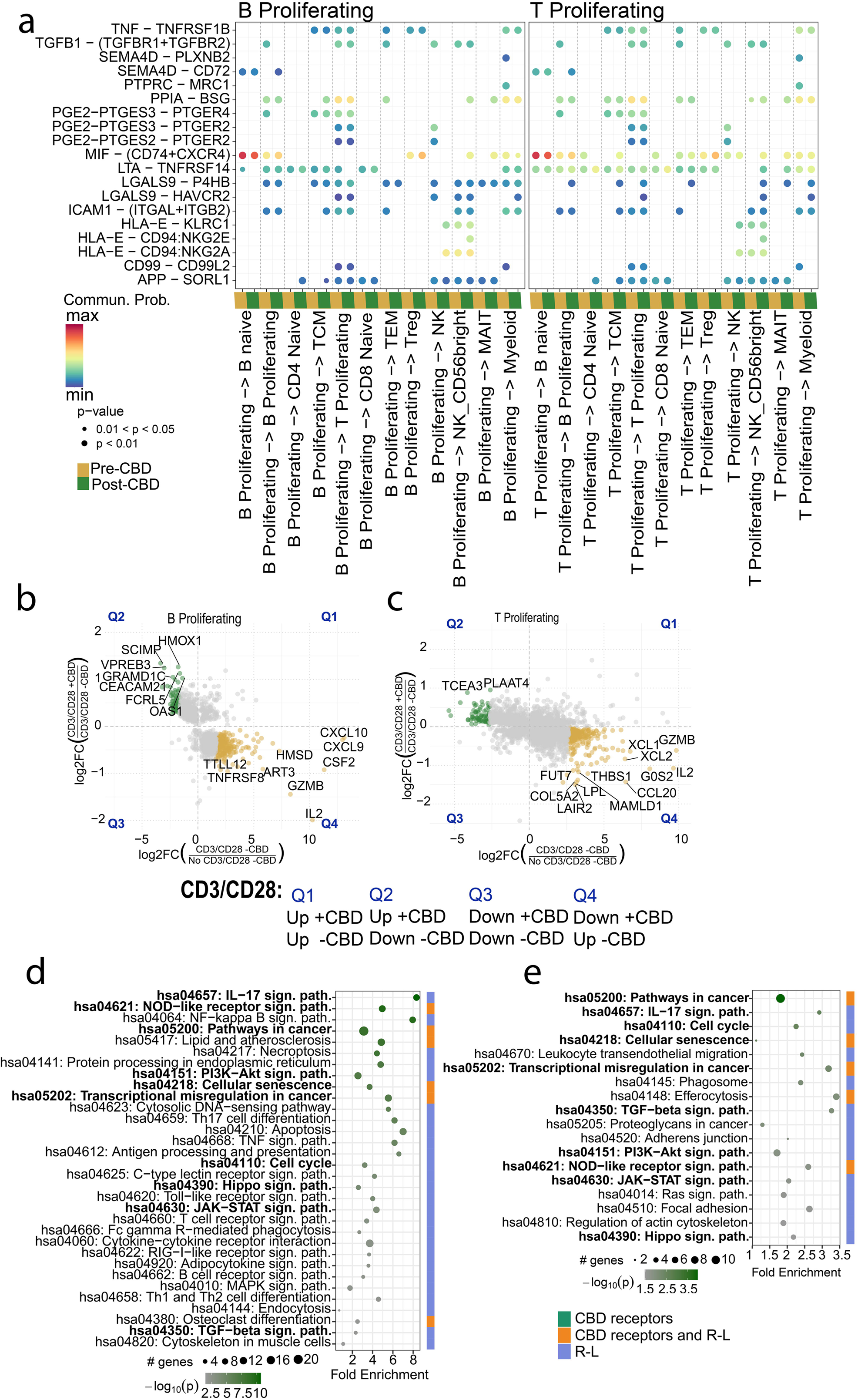
Proliferating B and T cell types. The receptor-ligand interactions, expression signature, and pathways of proliferating B and T cells were assessed pre-CBD and post-CBD. (a) Proliferating B and T cells were analyzed as the source population sending signals to other immune cells and themselves. R-L interactions that changed post-CBD included: TNF-TNFRSF1B, TGFB1 – (TGFBR1 + TGFBR2), and ICAM1 – (ITGAL+ITGB2). Receptor–ligand (R–L) communication profiles were inferred using CellChat for cells pre- and post-CBD treatment. The bubble plots show directional ligand–receptor (L–R) signaling where post-CBD has inhibition or activation. (b-c) Gene expression patterns in proliferating B (b) and T (c) cells exhibited similarities across two comparisons: *No CD3/CD28 -CBD vs CD3/CD28 -CBD* (x-axis) and *CD3/CD28 +CBD vs CD3/CD28 -CBD* (y-axis). The scatter plot depicts the log₂ fold changes (log₂FC) in gene expression. Each point represents a gene with statistically significant differential expression (adjusted *P-value* < 0.05) in both comparisons. Genes exhibiting opposing directions of regulation—up in one comparison and down in the other—are highlighted (green: upregulated with CBD under stimulation; gold: downregulated with CBD under stimulation). (d-e) The top 300 genes were ranked by Euclidean distance from the origin and used for pathway analyses for proliferating B cells (d) and T cells (**e**) via *pathfindR*. Genes with FDR-adjusted P-value < 0.05 were input into *pathfindR* using the BioGRID protein-protein interaction network and KEGG gene set. Enriched pathways were subsequently filtered to retain only those containing known cannabinoid (CBD) receptors or receptor–ligand (R–L) interactions. The enrichment chart displays the filtered pathways, ranked by –log₁₀(*P*), representing those most plausibly linked to CBD’s immunomodulatory actions.

B lymphocytes are known to express the cannabinoid receptor 2 on their cell surface (CB2 or *CNR2*), and reduced CB2 expression is associated with reduced cell proliferation^18^. Commensurate with this observation, we found greater expression of *CNR2* in proliferating B cells than naïve B cells. Further, the expression of *CNR2* was reduced post-CBD in all conditions, again consistent with CBD’s antiproliferative effects.

### PHARMACODYNAMIC INTERACTIONS OF CANNABIDIOL AND TACROLIMUS

The pharmacokinetic interaction between tacrolimus and CBD has been reported^5^. We sought to determine whether a pharmacodynamic interaction was also present between these drugs. Study participants took 11 days of oral cannabidiol; in contrast, tacrolimus (5 ng/ml) was administered *ex vivo* to the CD3/CD28 stimulated lymphocytes of each participant (for 48 hours) to avoid concentration variability resultant from the known pharmacokinetic interaction *in vivo*.

As a positive control, we confirmed that tacrolimus reduced cell proliferation (*P-value < 0.0001*, **Figure 6a**). We found that the reduction in proliferation was additive to that of CBD (*P-value < 0.01*, **Figure 6b**). We expanded the scRNA-seq object to 459,144 cells by adding the CD3/CD28 stimulated + tacrolimus conditions pre- and post-CBD. The absolute distribution of cells varied across the six conditions [No CD3/CD28, CD3/CD28, CD3/CD28 with tacrolimus; each pre- and post-CBD (**Figure 6c**)]. As expected, proliferating B and T lymphocytes decreased in the tacrolimus conditions (**Figure 6d**). Although there were substantial differences in cell type distribution, the expression patterns and cell cluster annotations were similar after tacrolimus addition (**Figure 6e**, **Supplementary Table 14**). As expected, tacrolimus administration proportionally reduced lymphocyte proliferation, with a relative increase in naïve B and T lymphocytes, both pre- and post-CBD (**Figure 6f**). The effects of CBD on TEM and TCM proportions were maintained even after tacrolimus addition (**Figure 6f**, **Supplementary Table 15**). Across all six conditions, we annotated unsupervised DEGs, supervised marker genes, CBD receptor expression, and cytokine expression. Genes associated with proliferation (*DLGAP5, KIF18B, KIF20A, MKI67, PBK*) were up-regulated after CD3/CD28, but down-regulated post-CBD (**Supplementary** Figure 3). The expression of *FABP5*, a purported CBD receptor, was decreased in the post-CBD conditions and with tacrolimus. Likewise, tacrolimus and CBD both prevented upregulation of transcript expression of the cytokines *CSF2*, *IL2*, *LTA*, and *TNF* (**Supplementary** Figure 3).

**Figure 6:**
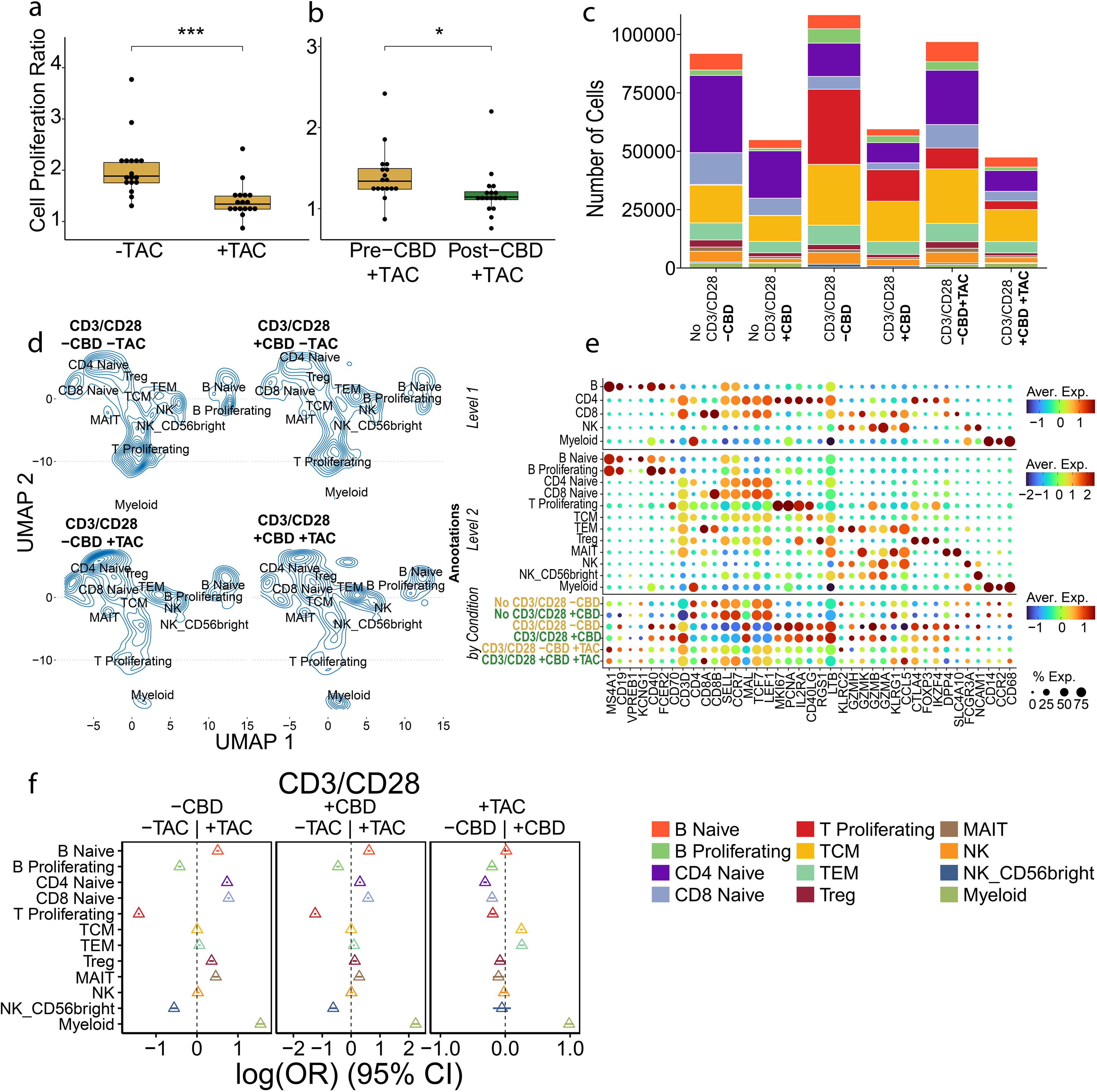
Pharmacodynamic interactions between cannabidiol and tacrolimus. The immune interactions between cannabidiol (CBD) and tacrolimus (TAC) were explored. Cells were exposed to CBD *in vivo* for 11 days, whereas TAC was added to cells *ex vivo*. (a) Cell proliferation was assessed by the CellTiter-Glo® assay in 2 conditions: CD3/CD28 -TAC (N=17) and CD3/CD28 +TAC (N=17). As expected, CD3/CD28 +TAC had reduced proliferation in the absence of cannabidiol compared to CD3/CD28 -TAC (*P-value < 0.0001, t-test*). (b) The effect of TAC on proliferation with and without CBD was compared in the pre-CBD (N=17) and post-CBD (N=18) timepoints. The relative proliferation of post-CBD +TAC was reduced compared to the pre-CBD +TAC condition (*P-value < 0.01, t-test*). All CD3/CD28 conditions were normalized to the No CD3/CD28 conditions to determine the change in proliferation in response to CD3/CD28. (c) Quantitation of absolute cell number in each scRNA-seq condition with/without CBD and TAC, split by Level 2 annotation. Number of cells (No CD3/CD28 -CBD -TAC= 91,865; No CD3/CD28 +CBD -TAC=54,914; CD3/CD28 -CBD -TAC= 108,423; CD3/CD28 +CBD -TAC=59,560; CD3/CD28 -CBD +TAC=96,914; and CD3/CD28 +CBD +TAC=47,468). (d) scRNA-seq cell density plot across conditions with/without CBD and TAC in the stimulated conditions. (e) Marker gene expression comparisons at different annotation resolutions and across the six conditions. A dot plot shows the average expression and cell percentage of pan marker genes for the PBMCs, B, T, NK, and myeloid cells. (f) The cell type distribution is expressed as an odds ratio for the Level 2 annotation and displayed in a forest plot for three comparisons. Left: Pre-CBD -TAC versus pre-CBD +TAC. Center: CD3/CD28 post-CBD -TAC versus CD3/CD28 post-CBD +TAC. Right: CD3/CD28 pre-CBD +TAC versus CD3/CD28 post-CBD +TAC. For each cell type, the central triangle marks the estimated difference in proportions between conditions, and the adjoining horizontal line depicts its 95 % confidence interval. A confidence interval that intersects the vertical dashed null line indicates no statistically significant difference, whereas intervals that remain entirely on one side denote significance at *P* -value< 0.05.

### CYTOKINE PROTEIN RESPONSE TO CBD

To orthogonally support the gene expression changes with functional protein secretion, we assessed *ex vivo* protein cytokine levels secreted from the lymphocytes, obtained in the same six conditions used in the scRNA- seq analysis. Based on a panel of protein cytokines, we observed a mixed pro-inflammatory and anti-inflammatory response in the post-CBD condition (**Figure 7** and **Supplementary** Figure 4, **Supplementary Table 16**). For example, we observed reduced TNF (**Figure 7a**) and LTA (**Figure 7b**) protein expression in the post-CBD condition, consistent with the reduced communication observed in TEM cells in the receptor-ligand analysis (**Figure 4a-b**). Likewise, proliferative signals were reduced in the post-CBD participants, including GM-CSF, IL-4, and IL-2 (**Figure 7c-e**). Specific anti-inflammatory cytokines, such as IL-10 and IL-1RA (antagonists of IL-1), were upregulated after CBD use (**Figure 7f and g**). In contrast, inflammatory cytokines (IL-8, CCL2, and IL-9) (**Figure 7h-j**) increased after CBD usage in the no CD3/CD28 and CD3/CD28 conditions. IL-6 (**Figure 7j**) was the most prominent inflammatory cytokine up-regulated in the post-CBD participants. Overall, the changes in all effector responses between no CD3/CD28 and CD3/CD28 were less potent (reduced delta) post-CBD, including innate (LTA or IL-12p40), intermediary innate-adaptive Th1 (TNF), adaptive (IL2), or Th2 (IL4). In contrast, the delta change of IL-10 is increased post-CBD, consistent with the relative increase in Treg population (for the CD3/CD28 condition, **Figure 2d**) and *KLF2*+ T cells (**Figure 4e**).

**Figure 7:**
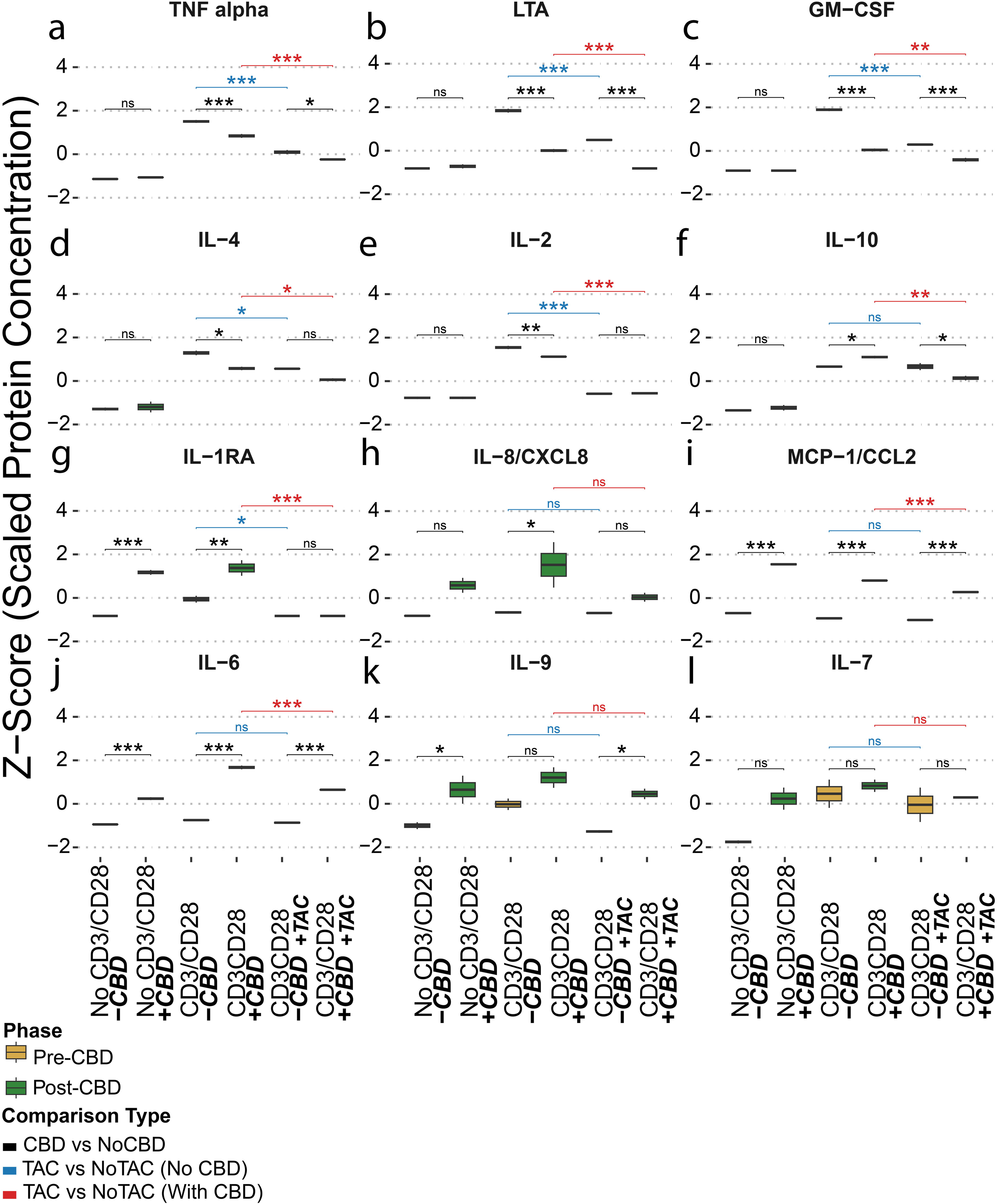
Cytokine levels. The effects of cannabidiol (CBD) treatment and/or tacrolimus (TAC) exposure on cytokine and chemokine levels were assessed under different conditions. CBD was administered *in vivo* over 11 days while TAC exposure was *ex vivo*. Each panel represents a box plot comparing pre-CBD (yellow) and post-CBD (green) conditions stratified according to No CD3/CD28 (No CD3/CD28), CD3/CD28 alone (CD3/CD28), and CD3/CD28 with Tacrolimus (CD3/CD28 + TAC). The same three participants provided samples for all six conditions. Due to the volume needed for the assay, biological samples were pooled, and technical replicates were used for statistics. Protein concentrations (pg/ml) were scaled and shown as z-score normalized values. Symbols indicate statistical comparisons: black denotes differences between pre- and post-CBD conditions; blue indicates comparisons between CD3/CD28 –TAC and CD3/CD28 +TAC in the absence of CBD; and red indicates the same comparison in the presence of CBD. Statistical significance is represented as follows: P-value< 0.05 (*), P-value < 0.01 (**), P-value < 0.001 (***), and “ns” for not significant. All P-values were calculated using the limma package with Benjamini–Hochberg false discovery rate (FDR) correction. Analytes shown include (a) TNF, (b) LT alpha (LTA), (c) GM-CSF, (d) IL-4, (e) IL-2, (f) IL-10, (g) IL-1RA, (h) IL-8/CXCL8, (i) MCP-1/CCL2, (j) IL-6, (l) IL-9, and (m) IL-7.

We next examined the effects of tacrolimus in addition to the CD3/CD28 condition with and without CBD to serve as a positive control (tacrolimus alone) and understand the combined effects of CBD and tacrolimus on cytokine production. Most cytokine expression was reduced following tacrolimus exposure. We observed an expected reduction in proliferative signals such as IL-2, GM-CSF, TNF, and LTA in both the CD3/CD28 pre-CBD and post-CBD conditions. We also observed a decrease in IL-10 post-CBD, but CCL2, IL-6, and IL-8 were still upregulated after CBD administration, even with combined tacrolimus exposure (**Figure 7**).

### FLOW CYTOMETRY VALIDATION OF LYMPHOCYTE POPULATIONS

To confirm the TEM/TEMRA cell upregulation seen in the scRNA-seq dataset by protein markers, we characterized the T cell distribution pre- and post-CBD with flow cytometry (N=3 participants, **Figure 8**). Cells were first gated as live/dead-CD45+ (viable leukocytes), followed by singlet selection using FSC-H vs FSC-A to exclude doublets. Live single lymphocytes were further classified into CD3+CD4+ (CD4 T cells) and CD3+CD8+ (CD8 T cells). Gating of CD8+ and CD4+ TEMs was established using CD8 or CD4 and a combination of CD45RO, CD45RA, and other markers (**Supplementary** Figure 5). The TEM cell distribution by flow cytometry mirrored that of the scRNA-seq datasets. CD4+ TEM cells (CD45RO+ CD45RA-) were upregulated post-CBD in the no CD3/CD28 and CD3/CD28 conditions. CD8+ TEM and TEMRA (CD45RO-CD45RA+) cells were increased in frequency post-CBD in most conditions (no CD3/CD28, CD3/CD28, and CD3/CD28+TAC, **Figure 8c-d**). We also identified a concomitant reduction in CD4 lymphocytes, consistent with the reduced proliferation in the scRNA-seq dataset (**Supplementary Table 17**). With tacrolimus administration, we identified expected changes in cell type distribution (**Supplementary Table 17**). However, CD8+ TEM and TEMRA cells were still relatively more abundant in the tacrolimus post-CBD condition compared to the tacrolimus pre-CBD condition (both *P-values* < 0.05).

**Figure 8:**
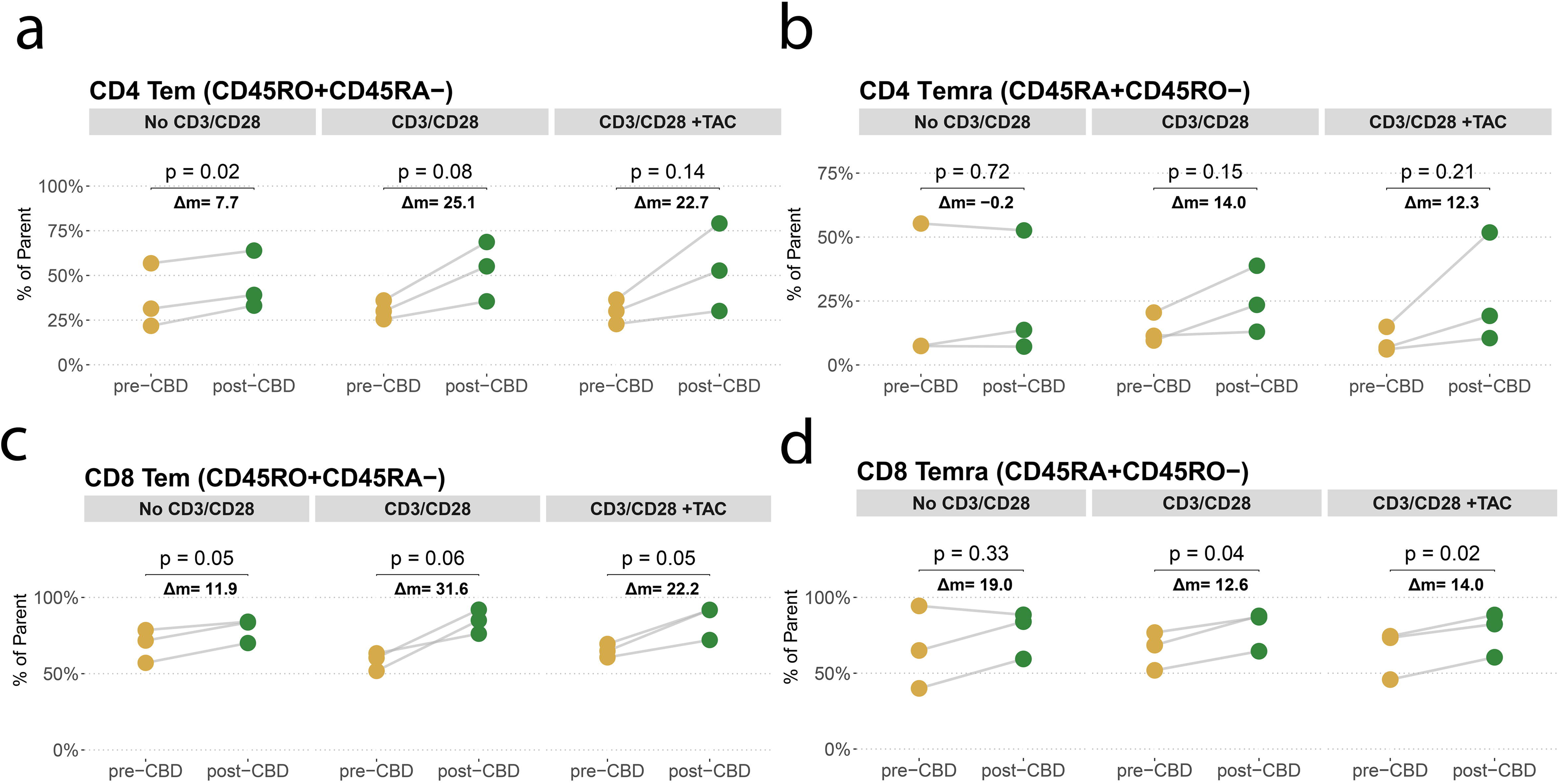
Flow cytometry of T Effector Memory Cells. Flow cytometric analyses of T lymphocyte cell subsets were conducted with corresponding statistical comparisons. Flow cytometry data for N = 3 participants illustrates the changes in cell type proportion pre- and post-CBD. Identified cells include a) CD4^+^ CD45RO⁺CD45RA⁻ (T effector memory, TEM), b) CD4^+^ CD45RA⁺CD45RO⁻ (TEMRA), c) CD8⁺ TEM, and d) CD8^+^ TEMRA T cell populations, each under pre- and post-CBD conditions across three experimental groups: No CD3/CD28 (No CD3/CD28), CD3/CD28 alone (CD3/CD28), and CD3/CD28 with Tacrolimus (CD3/CD28 + TAC). Significance determined by a paired t-test.

## Discussion

Cannabidiol (CBD) has gained widespread popularity and is now readily available without a prescription, raising important questions regarding its potential immunomodulatory effects in vulnerable populations such as allograft recipients. CBD is often regarded as safe due to its lack of psychoactive and addictive properties^19^; however, its FDA drug label includes warnings for hepatotoxicity and an increased risk of infections^20^. Prior clinical trials have examined the effect of CBD in human populations with multiple sclerosis^21,22^, revealing improvement in spasticity or other related neurologic symptoms. However, mechanistic insights were not proffered as to whether CBD’s immunomodulatory or neuro-modulatory effects drove these improvements.

Clinical studies addressing the CBD-induced immune changes in humans are still lacking. Many studies in animal or cell culture model systems have shown marked immunosuppressive effects of CBD^11,23–26^. For example, one study examined the immunosuppressive effects on peripheral blood mononuclear cells (PBMCs) of individuals with psoriasis vulgaris^27^, but a key difference is that CBD was only administered *in vitro* to cultured cells from these individuals. In contrast, our study is a clinical trial in which participants received oral CBD twice daily to reach a steady state, better reflecting the effects of the *in vivo* drug.

Our study demonstrates that CBD, when administered to human participants under steady-state conditions, exerts concentration-dependent immunomodulatory effects. We deeply profiled immune cell populations before (pre-CBD) and at steady-state (post-CBD) exposure using single-cell RNA sequencing. A key finding was a significant shift in the proportion of effector memory T cells, particularly the TEMRA phenotype, post-CBD. This shift is likely multifactorial. One factor is enhanced regulatory activity as supported by increased IL-10 cytokine expression, increased Treg to TEM ligand-receptor signaling through *LCK* and *HLA* genes/proteins, and a modest increase in the *FOXP3*+ Treg population in the scRNA-seq dataset. Prior studies have also demonstrated that CBD induces T regulatory lymphocytes^28^. In our study, the upregulation of *KLF2*+ T cells may similarly contribute to a quiescent T cell and/or TEMRA phenotype.^17,29^ A second potential factor is increased TEMRA/TEM terminal differentiation, potentially through the up-regulated IL-6. The cytokine IL-6 can promote the formation of effector cells that subsequently become TEM^30^, and CBD is known to stimulate IL-6 release^31^. Our clinical study confirmed the up-regulation of IL-6 secretion and flow cytometric increases in TEMRA/TEMs post-CBD. A third factor in the relative TEMRA/TEM upregulation is reduced proliferation and reduced total cell count of all cells. We observed a relative decrease in proliferating B and T cells through the anti-proliferative and cytotoxic effects of CBD. In contrast, TEMRA cells may persist longer under CBD conditions. TEMRA cells must first move to RO status, then express the death receptor (e.g. FAS), then move to an IL2Ra or subunit rearrangement status to increase their effector function before dying. The upregulated Treg activity may slow this process relative to the reduced proliferation of other T cells. Future mechanistic studies will be required to ascertain the realtive contribution of each of these factors and others to the TEMRA/TEM shift post-CBD. In summary, the results suggest that CBD influences T cell differentiation and memory formation, altering the immune balance.

CBD exposure reduced the proliferation of both T and B cells, consistent with prior reports of CBD’s anti-proliferative effects^10,11,32^. The effect is potentially secondary to the downregulation of cytokines and receptors such as IL-2 for T lymphocytes and IL-4 and CB2 for B lymphocytes^18,33^. In our study, *CNR2* (CB2) gene expression was reduced post-CBD. CB2 receptors are pre-proliferative and upregulated during inflammation, so *CNR2* down-regulation is consistent with an anti-inflammatory effect post-CBD.^34^ A pre-specified goal of this study was to deteremine whether CBD and tacrolimus held additive or synergistic immunologic effects. IL-2 protein and *IL-2* gene expression were suppressed by CBD, aligning with the known effects of tacrolimus, which also reduced IL-2 production. This suggests that CBD may share mechanistic overlap with tacrolimus, raising essential considerations for its use in allograft recipients or other immunocompromised populations.

The relative decrease in proliferating B and T cells, with increased proportions of TEMRA/TEMs, suggests mixed inflammatory and anti-inflammatory effects. Thus, our study demonstrates that CBD alters immune cell distribution and signaling in a nuanced and concentration-dependent manner.

In our assays, we selected against myeloid cells to better study CBD’s effect on lymphocyte populations. While we note a relative increase in myeloid cells in participants post-CBD, only a small number of myeloid cells were retained in the scRNA-seq populations (8,370 cells across all conditions compared to 450,774 lymphocytes).

Thus, we refrain from conclusions regarding the effects of CBD on myeloid cells given our experimental design that included positive lymphocyte selection. Prior investigations have focused on myeloid cells, identifying expression and phenotype changes. For example, certain studies have found that CBD induces an anti-inflammatory macrophage phenotype^26,27^. Other studies have shown that CBD induces apoptosis in macrophages^35,36^. We also identified a second clinical study that administered CBD at a daily dose of ∼1 mg/kg/day and identified an anti-inflammatory effect of CBD in their myeloid clusters (2,277 total myeloid cells)^37^.

Our study had certain limitations. First, this is a secondary analysis of individuals who participated in a pharmacokinetic clinical study^5^. As such, we were limited in the blood and sampling that could be performed outside the PK timepoints. Further, all single-cell power calculations were performed post-hoc, but the large quantity of cells (459,144) ensured adequate power to detect shifts in lymphocyte distribution. TEM and TEMRA cells are challenging to distinguish within scRNA-seq data; thus, we performed cytokine and flow cytometry analyses in a subset of participants to support the findings on an orthogonal protein level. Due to the prospective nature of this study, we cannot reinterrogate all individuals who previously completed the trial using every methodology. Finally, the dose of CBD administered (10 mg/kg/day total dose) is an FDA-approved dose, but higher than typical over-the-counter products (range: 0.1–5.6 mg/kg/day)^38^.

In conclusion, CBD appears to hold mixed pro- and anti-inflammatory effects. We believe the data presented here will be informative to transplant physicians who encounter patients who take CBD. It is now well known that CBD and tacrolimus interact pharmacokinetically, which can lead to unsafe elevations in tacrolimus concentration^4^. However, even when tacrolimus levels are fixed at 5 ng/ml, we have demonstrated important changes in cytokine signaling and immune cell distribution. These may hold downstream ramifications for preventing allograft rejection and resistance to opportunistic infections. Inflammatory IL-6 cytokine secretion and TEMRA/TEM cell proportion were upregulated in the post-CBD condition, regardless of the presence of tacrolimus. In particular, the changes in TEM proportion were concentration-dependent, suggesting providers must be aware of the total daily dose of CBD their patients take. Future studies must examine long-term outcomes in transplant recipients.

## Methods

### STUDY DESIGN AND PARTICIPANTS

The study is a secondary analysis of mechanistic molecular data acquired from healthy human volunteers who completed a phase 1 clinical study (NCT05490511) under Institutional Review Board (IRB) approval (IRB # 12763). Detailed descriptions of the trial design, eligibility criteria, interventions, and clinical procedures have been described^5^. Participants (N = 23) represent a pre-defined subset of the study population. Individuals were aged 18-65 without medical comorbidities or predicted drug-drug interactions (**Supplementary Table 1**). Blood samples were acquired at two time points. The first sample was acquired before the first CBD dose (pre-CBD). The second sample was acquired before the morning CBD dose on day 12 after 11 days of CBD titration (post-CBD). Most participants received EPIDIOLEX® (cannabidiol oral solution, GW Pharmaceuticals) at a dose of 2.5 mg/kg twice daily for 3 days, followed by 5 mg/kg twice daily for 11 additional days (14 total days), which is within the range of titration approved by the FDA. After 11 days of CBD administration, the participants underwent pharmacokinetic sampling at the following time points: 0h, 0.33h, 0.66h, 1h, 2h, 4h, 6h, 12h, 24h, and 48h.

### BLOOD SAMPLE PROCESSING

The peripheral blood mononuclear cell (PBMC) were acquired on day 1 (pre-CBD) and day 12 (post-CBD) from participants and then underwent 72 hours of ex vivo preparation for the following 6 conditions involving CBD, tacrolimus (TAC) and CD3/CD28 stimulation (CD3/CD28): 1) No CD3/CD28 / -CBD / -TAC, 2) CD3/CD28 / -CBD / -TAC, 3) CD3/CD28 / -CBD / +TAC, 4) No CD3/CD28 / +CBD / -TAC, 5) CD3/CD28 / +CBD / -TAC, 6) CD3/CD28 / +CBD / +TAC. Blood samples were diluted 3-fold using Dulbecco’s Phosphate-Buffered Saline (DPBS) [Corning Inc., Corning, NY; #21-031-CM] with 1% fetal bovine serum (FBS) [Gibco, Grand Island, NY; #A3840101] and added atop of 15 mL of Ficoll-PaqueTM PLUS (Cytivia, Marlborough, MA; #17144002) in a tube, which was centrifuged at 600 g for 30 minutes at room temperature. The buffy coat containing PBMCs atop the Ficoll layer was pipetted into a tube and washed with DPBS with 1% FBS twice with repeated centrifuging and discarding of supernatant. Finally, the cell pellets were resuspended in 10 mL of RPMI 1640 medium (Roswell Park Memorial Institute, Corning Inc., Corning, NY; #10-040-CM) with 1% FBS and 1% penicillin-streptomycin (p/s) to which 50 μL of phytohemagglutinin-L (PHA-L) [Millipore-Sigma, Burlington, MA; #11249738001] was added for preferential lymphocyte selection. Each participant’s cells were transferred to a T25 flask and incubated at 37°C and 5% CO2 for 24 hours of *ex vivo* primary culture.

After 24 hours, floating cells not affixed to the T25 flask were harvested and centrifuged at 600 g for 5 minutes at room temperature. Subsequently, the supernatant was discarded, and the cells were resuspended in RPMI. Samples of each participant were divided into three conditions and incubated in T25 flasks or 96-well plates at 37°C and 5% CO2 for 48 hours. The first condition (CD3/CD28) received CD3/CD28 lymphocyte stimulation at 25 μL/mL (ImmunoCult™ Human CD3/CD28 T Cell Activator) [STEMCELL Technologies #10971]. The second condition (CD3/CD28 + TAC) received CD3/CD28 lymphocyte stimulation with tacrolimus at a concentration of 5 ng/mL [MedChemExpress; #HY-13756/CS-1507]. A third condition (No CD3/CD28 / baseline) did not have CD3/CD28 activators or tacrolimus added to the samples.

After 72 hours, cells were retrieved. Cells in T25 flasks were used for scRNA-seq, flow cytometry, or cytokine measurement. The cells were centrifuged at 600 g for 5 minutes at room temperature, washed, and resuspended in RPMI for downstream processing. Cells from 96-well plates were used for the cell proliferation assay.

### LUMINESCENCE ASSAY

Cell viability using CellTiter-Glo® [Promega Corporation, Madison, WI; CAT#G7570] was measured in cells on 96-well plates to quantify cell proliferation by adenosine triphosphate (ATP) detection using a Spectramax M5 system coupled with SoftMax Pro 5.2 (Molecular Devices, San Jose, CA). Pre-CBD (N = 17) and post-CBD (N = 18) values were calculated as a ratio of ATP detection in the stimulated (CD3/CD28) condition normalized by the non-stimulated (No CD3/CD28) condition. Ratios were separately calculated with and without tacrolimus. Conditions were compared using a t-test (**Supplementary Table 2**).

### PBMC SINGLE-CELL RNA SEQUENCING (scRNA-seq)

For each of the six conditions in **Supplementary Table 1**, library preparation was performed using the 10x Genomics Chromium Single Cell V3.1 NextGem platform. cDNA synthesis and amplification were performed according to the 10x Genomics protocols to ensure optimal yield and specificity. Quality control measures were applied at multiple stages to assess library integrity, concentration, and fragment size; sequencing was conducted on the Illumina NovaSeq 6000 platform. Paired-end reads were generated with a length of 100 base pairs (PE100), and sequencing depth was optimized according to experimental requirements. Fifthy nine scRNA-seq experiments were sequenced across multiple runs to achieve sufficient coverage for downstream analyses (**Supplementary Table 1**). Data were obtained from 12 participants who provided pre-CBD scRNA-seq samples, eight of whom also offered post-CBD scRNA-seq samples, which passed QC. Two participants withdrew before the post-CBD sample acquisition, and two samples did not pass QC. Alignment and quantification were accomplished with CellRanger (v. 7.1.0) using default options and reference genome GRCh38-2020-A. The BPCells (v.0.2.0) package converted the matrices in the Hierarchical Data Format h5 to bitpacked compressed format stored as binary files on a hard drive. We used Seurat (v.5.1.0) to create a Seurat object for each sequencing sample. Cells with mitochondria reading over 25% or fewer than 500 genes were removed. All scRNA-seq samples were merged, and the cells were annotated by Azimuth (v.0.5.0) using the PBMC_ref_4546839 in Zenodo (https://zenodo.org/records/4546839).^39^ For the six conditions, the scRNA-seq object was split into layers by sample, with the following methods: NormalizeData, FindVariableFeatures, ScaleData, and IntegrateLayers, finally using the method HarmonyIntegration with the harmony (v.1.2.0) tool performed to integrate the samples.

Two additional participants provided snRNA-seq at the post-CBD condition, which only increased the sample size for comparisons between pharmacokinetics and cell type proportion (**Figure 3**). These samples were analyzed separately and not included in the Seurat object or other scRNA-seq analyses beyond those of **Figure 3**. Cell annotations were performed in an analogous way to that of scRNA-seq. A principal component analysis for cell type proportion did not reveal differential clustering of the scRNA-seq and snRNA-seq samples.

### CELL PROPORTION QUANTIFICATION

The proportions of each cell type across conditions were calculated to assess differences in cell distribution. Cells in each condition were grouped into cell type, and the count of each cell type was divided by the total cell count within that condition to determine the cell type-specific proportions. Odds ratios were calculated by comparing the proportions between conditions (e.g., A and B) for each cell type, specifically by computing:

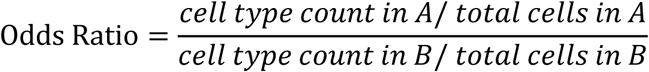

Significance of the observed difference was evaluated using a Fisher’s exact test, with a P-value < 0.05 to indicate significance.

Confidence intervals (95% CI) for the odds ratios were calculated by taking the natural logarithm of the odds ratio and applying the following formula:

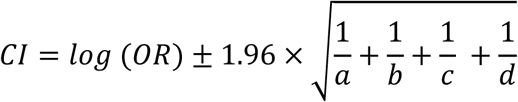

Where a, b, c, and d represent the cell counts in each condition matrix for the analyzed cell type. The 95% CI was then exponentiated to obtain the final interval on the odds ratio scale, and results were visualized through a forest plot.

### PHARMACOKINETICS (PK) OF CBD AT STEADY STATE

CBD steady-state concentrations were obtained using an LC-MS/MS Assay^5^. Ten participants provided 10 PK time points with CBD concentrations. The data were analyzed by noncompartmental analysis using Phoenix® WinNonlin® version 8.4 (Certara L.P., Princeton, NJ). The concentration at time 0h (C₀_h_) in ng/mL, maximum blood concentration (C_max_) in ng/mL, time to C_max_ (T_max_) in hours (h), area under the plasma concentration-versus-time within a dosing interval or AUCTau (h*ng/mL), C_Tau_ (ng/mL), minimum concentration (C_min_) in ng/mL and time at C_min_ or T_min_ (h).

Descriptive statistics were computed for each parameter, including mean, median, minimum (Min), maximum (Max), standard deviation (SD), and coefficient of variation (CV%). Given the small sample size, a t-distribution was used to estimate the 95% confidence interval (CI). The confidence coefficient was determined using the t critical value for a 95% confidence level with 9 degrees of freedom. The margin of error (ME) was calculated by multiplying the t critical value by the standard error. The 95% confidence interval was computed as Mean ± Margin of Error, providing upper and lower limits.

### CORRELATION OF PK TO CELL PROPORTION

To assess the correlation between sc/snRNA-seq and pharmacokinetic (PK) parameters, we analyzed data from 10 participants (N=10) who had measurements available for both data types under the post-CBD (CD3/CD28 / +CBD / -TAC) condition. We applied linear regression using the lm() function in R to model the relationship between an independent variable (C_0h_) and the dependent variable (TEM proportion). The C_0h_ parameter was selected for comparison because it is the same concentration time point at which participants had their sc/snRNA-seq sample drawn. Pearson correlation was used to evaluate the strength of the association, with statistical significance at P-value < 0.05. The R-squared value from the regression indicates the proportion of variance in cell type proportion explained by C_0h_.

### IMMUNE CELL-CELL COMMUNICATION

Analysis of scRNA-seq datasets was performed by CellChatv2 (v.2.1.2)^40^ to assess the cell-cell communication across the cell type and compare the pre-CBD and post-CBD stimulated conditions (**Supplementary Table 6**).

### PATHWAY ANALYSES

Differentially expressed genes (DEGs) were identified across conditions of interest using the default settings of RunPresto from the SeuratWrappers (v0.3.5) and presto (1.0.0) packages^41^. DEGs were filtered to include only genes with a P-value by Wilcox and adjusted P-value < 0.05 after Benjamini-Hochberg correction (BH) and were ranked by the absolute value of log-fold change (*|avgLogFC|*). To identify the gene set enhanced and suppressed by CBD, a plot on the x-axis of DEGs from No CD3/CD28 -CBD versus CD3/CD28 -CBD, and the y-axis CD3/CD28 CBD versus CD3/CD28 -CBD. The genes in quadrant 2 (Q2) are enhanced by CBD, and the genes in quadrant 4 (Q4) are suppressed by CBD. The top 300 ranked by Euclidean distance from the origin are annotated and used for pathway enrichment analysis. Pathway enrichment analysis was conducted using pathfindR (v2.4.1).^42^ The enrichment parameters for run_pathfindR were configured with the KEGG gene sets and the Biogrid protein interaction network, allowing for pathways with a minimum of 1 and a maximum of 500 genes, and the adjusted P-value < 0.05. The enriched pathways were screened for the presence of genes in pathways related to CBD receptors^3^ and receptor-ligands identified in the cell-cell communication analyses (Supplementary Tables 8, 11, and 13).

### PBMC CYTOKINE MULTIPLEX ASSAY

Cells from the six conditions were used in a multiplex assay. The supernatant of lymphocytes was aliquoted in 1.5 mL tubes and kept at -80°C until use. Samples from 3 participants then underwent multiplex assays using ProcartaPlexTM Human Cytokine/Chemokine/Growth Factor Convenience Panel 1 (Invitrogen, Thermo Fisher Scientific Inc., MA, USA., #EPXR450-12171-901). The samples were pooled by condition, and comparisons were made between the pre-CBD and post-CBD time points (**Supplementary Table 1**).

For the multiplex assay, samples were thawed on ice and assayed by Luminex xMAP technology for 45 human cytokines and chemokines (**Supplementary Table 17**). Cytokine concentrations were quantified from culture supernatants with the 45-plex Luminex® assay and exported as pg mL⁻¹ values. The genes (cytokine) × samples (well) expression matrix were analyzed with limma (v 3.60.6). A no-intercept design matrix encoded every experimental condition: 1) No CD3/CD28 / -CBD / -TAC, 2) CD3/CD28 / -CBD / -TAC, 3) CD3/CD28 / -CBD / +TAC, 4) No CD3/CD28 / +CBD / -TAC, 5) CD3/CD28 / +CBD / -TAC, 6) CD3/CD28 / +CBD / +TAC.

Differential cytokine abundance was tested with limma’s moderated t-statistic, controlling for the false-discovery rate with a BH adjustment at FDR < 0.05.^43^

### PBMC FLOW CYTOMETRY

After 72 hours of primary culture, petri dishes containing both attached and floating cells were scraped, and the culture media containing the cells were centrifuged at 1600 × g for 5 minutes. The cell pellets were transferred to 2 mL tubes, and CryoStor®CS10 was added. The cells were placed on ice for 10 minutes, stored overnight in a deep freezer within a slow-freeze container, and finally moved to liquid nitrogen until used for flow cytometry analyses. Thawed cells were resuspended in fresh culture media (RPMI supplemented with 1% FBS and 1% penicillin/streptomycin), centrifuged (350 × *g*, 7 min, 4°C), and supernatants were aspirated. Cells were incubated with Live/Dead Olive dye (1:1000 dilution in dPBS) for 30 min at 4°C (light-protected), washed by centrifugation (350 × *g*, 7 min, 4°C), and supernatants were removed. Fc receptors were blocked with FcBlock (5 µL/tube, 10 min, 4°C). Antibody cocktails, single stains, and fluorescence-minus-one (FMO) controls were prepared and added to cell suspensions. After 30 min of incubation (4°C, light-protected), cells were washed with 2 mL flow cytometry staining buffer (PBS + 2% FBS) and centrifuged (350 × *g*, 7 min, 4°C).

For intracellular staining, cells were fixed and permeabilized using IC Fixation/Permeabilization Buffer (20 min, room temperature, light-protected), washed with 2 mL permeabilization buffer, and centrifuged (350 × *g*, 7 min, 4°C). Permeabilized cells were resuspended in 100 µL permeabilization buffer, incubated with intracellular antibody cocktails (20 min, room temperature, light-protected), rewashed with 2 mL permeabilization buffer, and finally resuspended in flow cytometry staining buffer for acquisition. Single antibody staining controls were prepared using Ultracomp eBead Plus. All samples were analyzed using the Cytek Aurora System. Flow cytometry population frequencies were normalized by rarefying subpopulation counts to the minimum CD3⁺ T cell count per condition. Differential abundance between conditions was assessed using two-sided Fisher’s Exact Tests on 2×2 contingency tables, followed by BH correction for multiple comparisons^44^.

For gating, live singlet lymphocytes were selected by forward scatter area and forward scatter height with exclusion based on the viability dye. Total T cells were identified as CD3⁺ events and split into CD4⁺ and CD8⁺ lineages. Within the CD8+ compartment we resolved memory/effector phenotypes on the basis of CD45 isoforms: CD45RO⁺ CD45RA⁻ effector-memory (TEM), CD45RA⁺ CD45RO⁻ (TEMRA), and an intermediate CD45RA⁺ CD45RO⁻ pool that was further subdivided into intermediary TEM, TCM (central memory) and naive/TEMRA subsets according to granzyme-A, TGF-β, CD127 and CD25 expression. CD4+ T cells were partitioned into regulatory (CD25^high^ FoxP3⁺) and non-T-regulatory (CD25^low^ FoxP3⁻) fractions; each fraction was then stratified in an identical CD45RO/CD45RA hierarchy yielding TEM, TCM, intermediary TEM, naive and TEMRA populations.

## ACKNOWLEDGMENTS

This work was supported by the National Institute of Health/National Center for Complementary and Integrative Health (NIH/NCCIH) (MTE, DLG, Grant R01AT011463) and the NIH/ National Institute of General Medical Sciences (NIH/NIGMS) (ZD, Grant R35GM145383). The Intramural Research Program of the National Institutes of Health partly supported this research. GCS and JE were/are supported by the National Institutes of Health and the National Institute of General Medical Sciences (Grant T32GM008425).

## AUTHOR CONTRIBUTIONS

D.L.G. and M.T.E. wrote the manuscript. D.L.G., A.A.S., Z.D., and M.T.E. designed the research. G.C.S., J.B.L.L., D.J.F., S.K., Y-H.C., K.M., J.E., T.R.B., Z.C., K.M., M.A., E.R., Y.H.C., J.S.S., R.B.P., Z.D., and M.T.E. performed the research. D.L.G, G.C.S., D.J.F., S.K., J.E., M.S.M., P.C.D., L.S., R.B.P., Z.D., and M.T.E. analyzed the data. M.T.E. contributed new reagents/analytical tools.

## COMPETING INTERESTS

The authors have nothing to disclose. MATERIALS & CORRESPONDENCE

Please address data and materials requests to Michael Eadon.

## Code Availability

Most code methods are derived from standard packages referenced in the methods. Full code will be made available through GitHub upon acceptance.

## Data availability

Data is available within the Gene Expression Omnibus (GEO) under GSE303581.

---

**Supplementary Figure 1:**
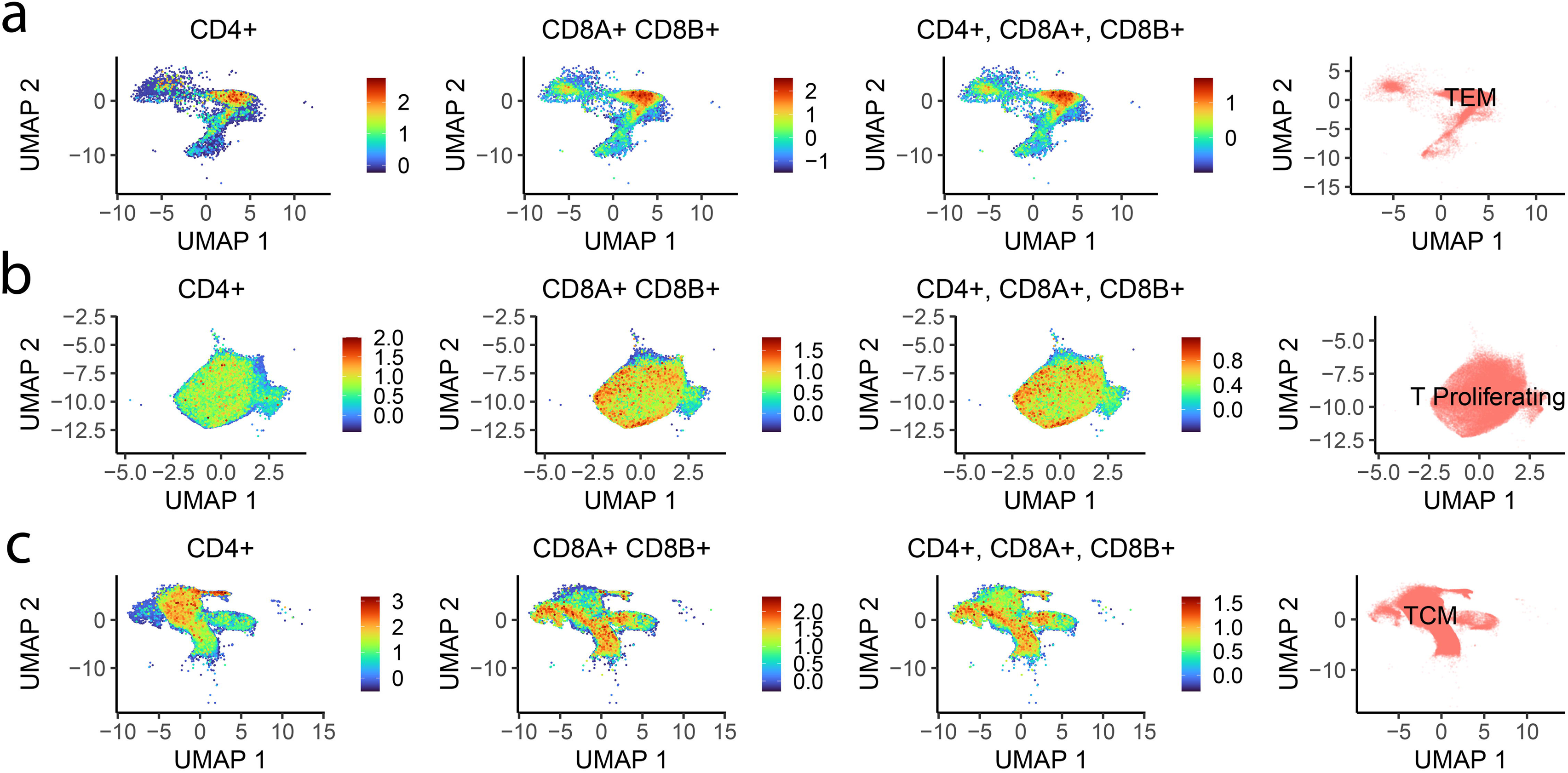
Phenotype-Driven Annotation of T-Cell Clusters. CD8+ and CD4+ T lymphocytes were sub-clustered to assess overlap of their populations based on transcriptional profiles. The UMAP visualization shows clustering of T cells, with annotations highlighting major phenotypic subsets. In some cases, CD8+ and CD4+ T cells are resolved into distinct clusters, while in others, they are grouped. Clustering was primarily driven by phenotypic features rather than strict lineage (CD8/CD4) identity, reflecting shared functional or transcriptional programs across subsets. The feature plots are showing this effect occurring in (a)T effector memory (TEM), (b) T Proliferating and (c) T central memory (TCM).

**Supplementary Figure 2:**
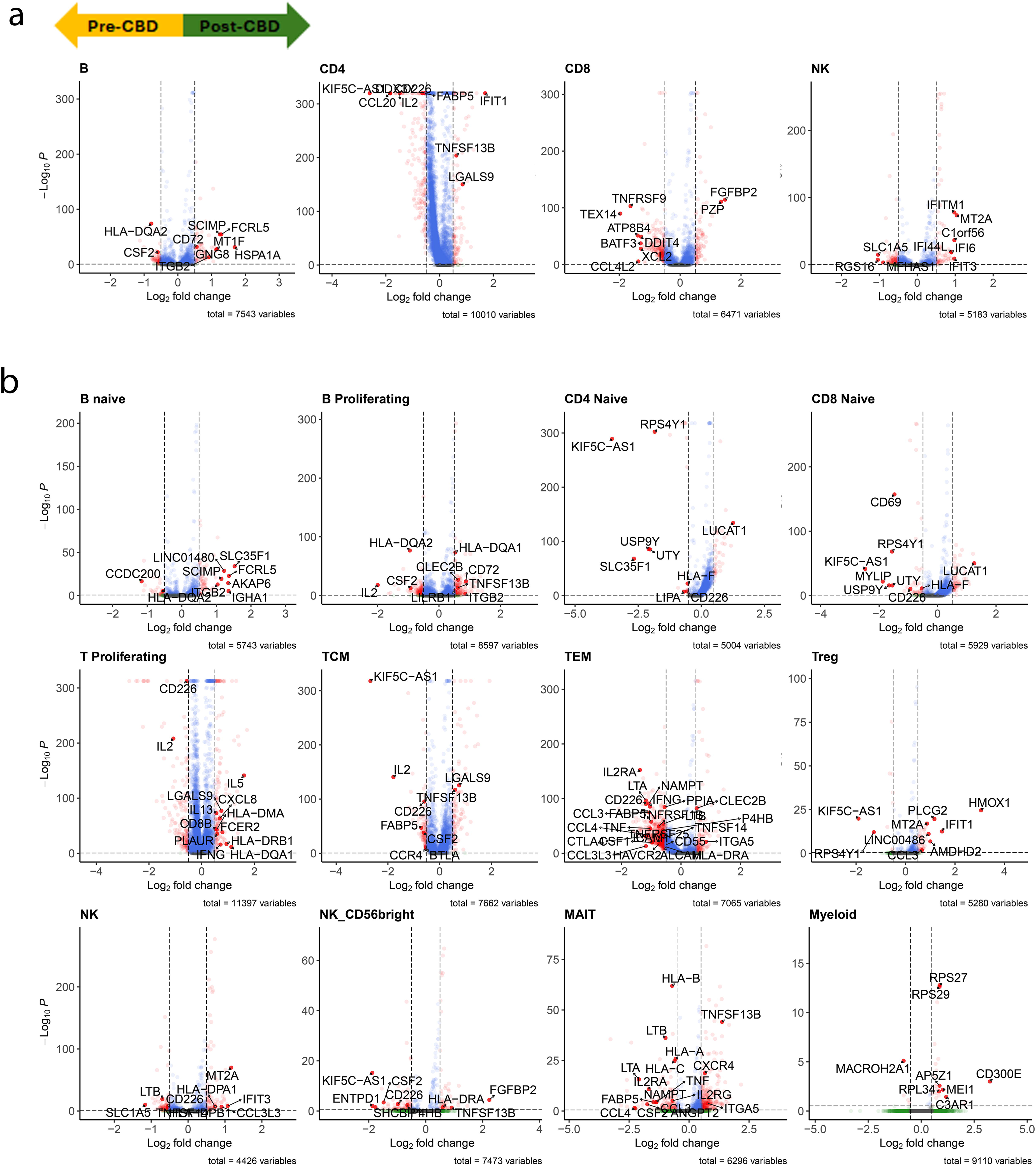
Differential expression genes (DEGs) in CD3/CD28 –CBD versus CD3/CD28 +CBD. The volcano plots show red dots for genes that have average logFC > 0.5 and an adjusted P-value < 0.05 (Wilcox and Benjamini-Hochberg). Blue dots correspond to adjusted average logFC < 0.5 and adjusted P-value < 0.05. Green genes correspond to an average logFC > 0.5 and adjusted P-value > 0.05. Grey dots correspond to an average logFC < 0.5 and adjusted P-value > 0.05. Negative x-axis are genes downregulated, CD3/CD28 post-CBD, and positive x-axis are upregulated CD3/CD28 post-CBD. (a) DEGs at annotation Level 1: B, CD4, CD8, NK and Myeloid (b) DEGs at annotation Level 2: B naive, B Proliferating, CD4 Naive, CD8 Naive, T Proliferating, T Central Memory (TCM), T effector memory (TEM), T regulatory (Treg), Mucosal-associated invariant T (MAIT), Natural Killer (NK), and Natural Killer CD56+ (NK_CD56bright).

**Supplementary Figure 3:**
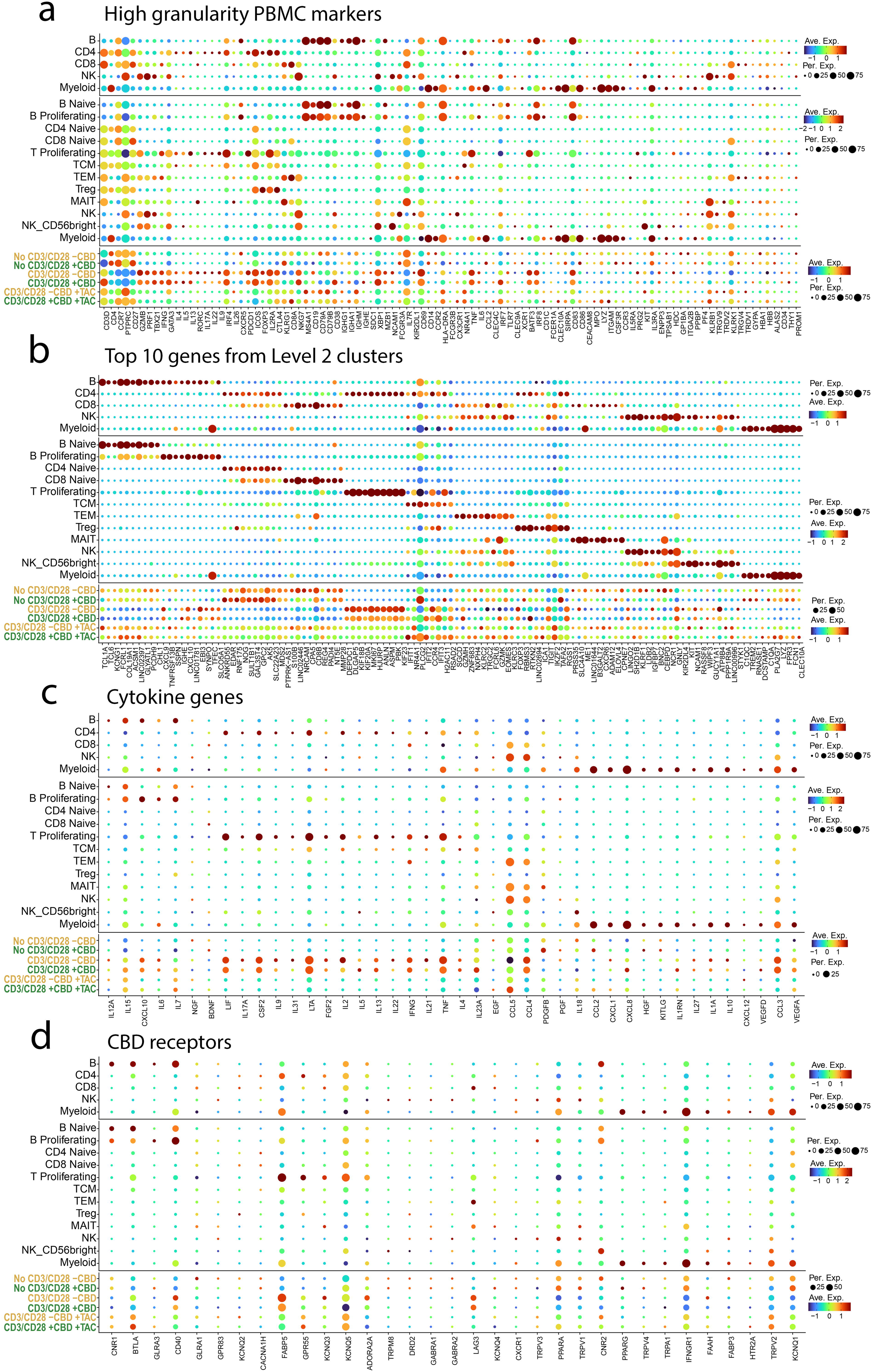
Comparative Gene-Set Dotplots Across Three Cell Annotations. The Dot plot shows the expression of the genes across 3 annotation levels: Level 1 (B, CD4, CD8, NK and Myeloid), Level 2 with 12 cell types (B Naïve B proliferating, CD4 Naïve, CD8 Naïve, T proliferating, TCM, TEM, Treg, MAIT, NK, NK_CD56bright and Myeloid), and Level 3 for the 6 conditions: No CD3/CD28 –CBD, No CD3/CD28 +CBD, CD3/CD28 -CBD, CD3/CD28 +CBD, CD3/CD28 –CBD +TAC and CD3/CD28 +CBD +TAC. (a) PBMC markers with high granularity to identify the main immune cell population. (b) Top 10 differentially expressed genes (DEGs, unbiased) for each cluster based on Level 2 annotation, but plotted for all three annotation levels. (c) cytokine proteins using the respective gene symbol. (d) CBD receptors were used in our analyses to filter the pathway analyses.

**Supplementary Figure 4:**
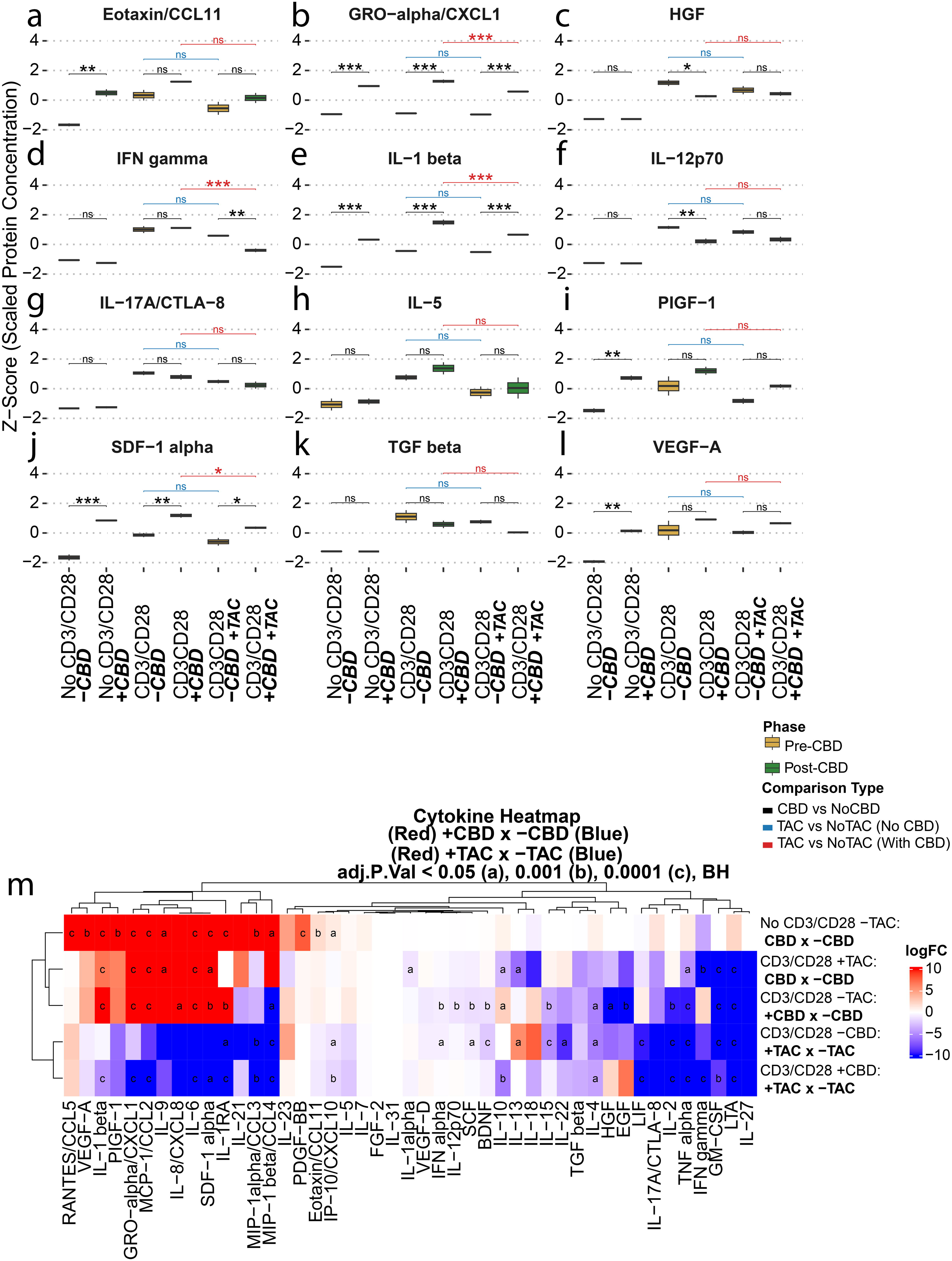
Additional cytokine and chemokine levels in the (±CD3/CD28), (±CBD), and (±TAC) conditions. Each panel presents box plots comparing cytokine levels in the pre-CBD (yellow) and post-CBD (green) conditions across three experimental groups: No Stimulation (No CD3/CD28), Stimulation alone (CD3/CD28), and Stimulation with Tacrolimus (CD3/CD28 + TAC). Protein concentrations are shown as z-score normalized values (pg/ml). Symbols indicate statistical comparisons: black denotes differences between pre- and post-CBD conditions; blue indicates comparisons between CD3/CD28–TAC and CD3/CD28 +TAC in the absence of CBD; and red indicates the same comparison in the presence of CBD. The statistical significance is represented as follows: P-value < 0.05 ***a***, P-value < 0.01 ***b***, P-value < 0.001 ***c***. All p-values were calculated using the limma package with Benjamini–Hochberg false discovery rate (FDR) correction. Analytes shown include: (a) EOTAXIN/CCL11, (b) GRO-alpha/CXCL1, (c) HGF, (d) IFN gamma, (e) IL-1 beta, (f) IL-12p70, (g) IL-17A/CTLA-8, (h) IL-5, (I) PGF1, (j) SDF-1, (k) TGF beta, and (l) VEGF-A. The heat map (m) provides an integrated overview of cytokine and chemokine responses across the same experimental comparisons depicted in the box plots. Each row corresponds to a specific group-wise comparison—namely, pre-CBD versus post-CBD, CD3/CD28 –TAC versus CD3/CD28 +TAC in the absence of CBD, and CD3/CD28 –TAC versus CD3/CD28 +TAC in the presence of CBD—aggregated across all measured cytokines. This visualization enables the identification of coordinated immunological patterns and potential treatment-related effects, offering a comprehensive, systems-level perspective on the modulatory impact of CBD and Tacrolimus.

**Supplementary Figure 5:**
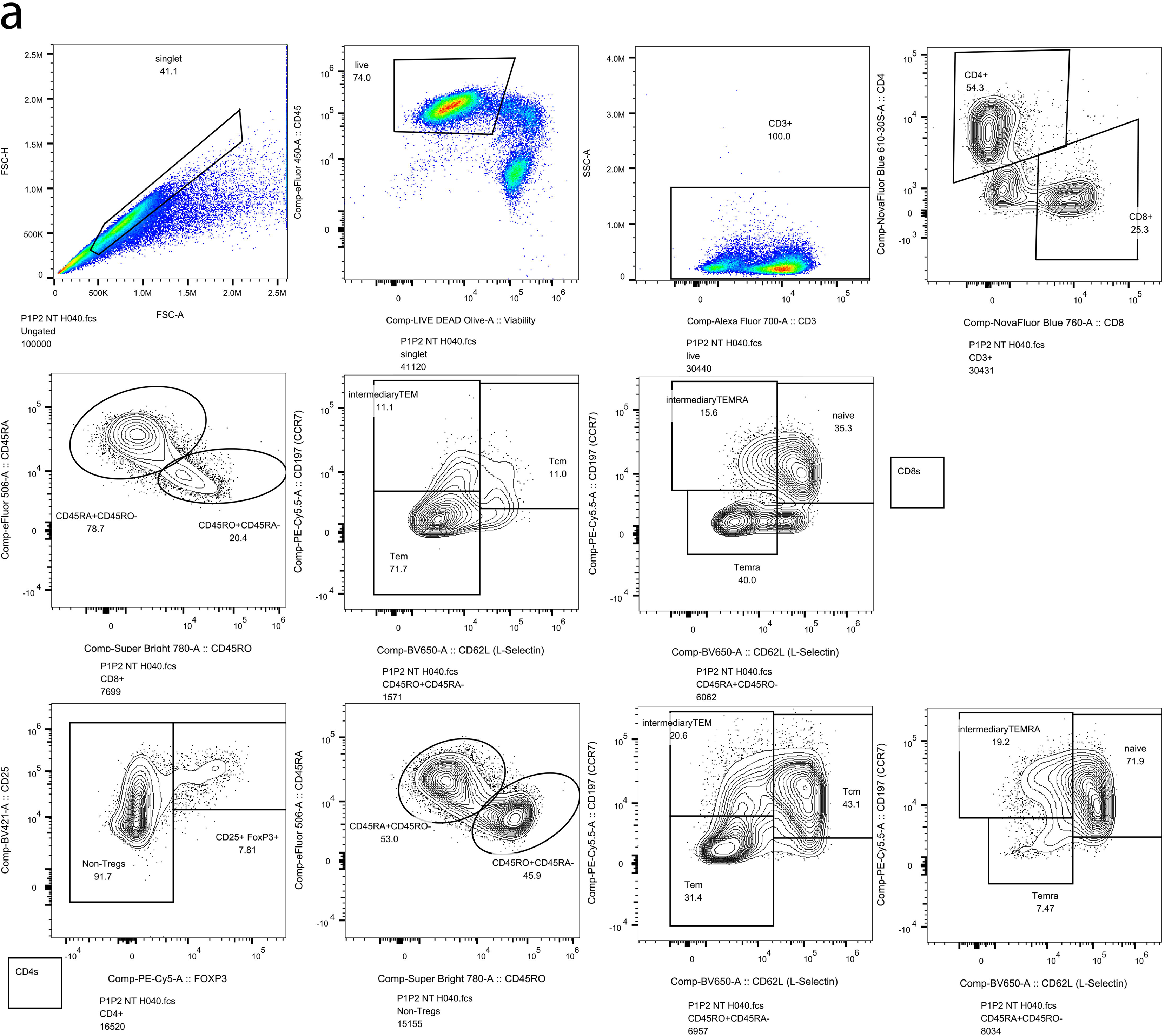
Representative flow-cytometry gating strategy used to delineate T-cell subsets. (a) Live singlet lymphocytes were first selected by FSC-A × FSC-H and exclusion of the viability dye. Total T cells were identified as CD3⁺ events and subsequently split into CD4⁺ and CD8⁺ lineages. Within the CD8 compartment we resolved memory/effector phenotypes on the basis of CD45 isoforms: CD45RO⁺ CD45RA⁻ effector-memory (Tem), CD45RA⁺ CD45RO⁻ (Temra), and an intermediate CD45RA⁺ CD45RO⁻ pool that was further subdivided into intermediary Tem, Tcm (central memory) and naive/Temra subsets according to granzyme-A, TGF-β, CD127 and CD25 expression. CD4 T cells were first partitioned into regulatory (CD25^high FoxP3⁺) and non-T-regulatory (CD25^low FoxP3⁻) fractions; each fraction was then stratified in an identical CD45RO/CD45RA hierarchy yielding Tem, Tcm, intermediary Tem, naive and Temra populations. Median fluorescence of CD127 (IL-7Rα) and CD25, as well as granzyme-A⁺ and TGF-β⁺ functional overlays, are displayed for the indicated gates. Percentages shown correspond to the frequency of each subset within its parent population. Abbreviations: Tem, effector memory; Temra, terminally differentiated effector memory re-expressing CD45RA; Tcm, central memory; cTreg, conventional regulatory T cell. The participant in the condition No CD3/CD28 -CBD -TAC (P1P2 NT H040) starts with 30440 CD3^+^ cells where 54.3% is CD4^+^ and 25.3% is CD8^+^.

## Supplementary Tables Title

Supplementary Table 1: Participants Stratified Across Experimental Conditions and Technologies Supplementary Table 2: CellTiter-Glo® data

Supplementary Table 3: Differentially Expressed Genes (DEGs) Across the Three Annotation Levels Supplementary Table 4: Cell Proportion Forest Plot pre-CBD versus post-CBD

Supplementary Table 5: Volcano Plots CD3/CD28 +CBD -TAC versus CD3/CD28 -CBD -TAC Supplementary Table 6: Cell Cell Communication Result Across Level 2 Annotation

Supplementary Table 7: Dual-Contrast CBD Modulation Analysis in Effector-Memory T (TEM) Supplementary Table 8: Pathway Analysis with KEGG for Effector-Memory T (TEM) cells

Supplementary Table 9: Network of CBD-Regulated Pathways in Effector-Memory T (TEM) cells

Supplementary Table 10: Dual-Contrast CBD Modulation Analysis in Proliferating B cells

Supplementary Table 11: Pathway Analysis with KEGG for Proliferating B cells

Supplementary Table 12: Dual-Contrast CBD Modulation Analysis in Proliferating T cells

Supplementary Table 13: Pathway Analysis with KEGG for Proliferating T cells

Supplementary Table 14: Differentially Expressed Genes (DEGs) Across the Three Annotation Levels Supplementary Table 15 | Cell Proportion Forest Plot with Tacrolimus

Supplementary Table 16 | Cytokine Data

Supplementary Table 17 | Flow Cytometry Cell Counts and Proportions

